# Protein kinase B controls *Mycobacterium tuberculosis* growth via phosphorylation of the global transcriptional regulator Lsr2

**DOI:** 10.1101/571406

**Authors:** Kawther Alqaseer, Obolbek Turapov, Philippe Barthe, Heena Jagatia, Angélique De Visch, Christian Roumestand, Malgorzata Wegrzyn, Iona L. Bartek, Martin I. Voskuil, Helen O’Hare, Adam A. Witney, Martin Cohen-Gonsaud, Simon J. Waddell, Galina V. Mukamolova

## Abstract

*Mycobacterium tuberculosis* is able to persist in the body through months of multi-drug therapy. Mycobacteria possess a wide range of regulatory proteins, including the essential protein kinase B (PknB), that control transitions between growth states. Here, we establish that depletion of PknB in replicating *M. tuberculosis* results in transcriptional adaptations that implicate the DNA-binding protein Lsr2 in coordinating these changes. We show that Lsr2 is phosphorylated by PknB, and that phosphorylation of Lsr2 at threonine 112 is important for *M. tuberculosis* growth and survival under hypoxic conditions. Fluorescence anisotropy and electrophoretic mobility shift assays demonstrate that phosphorylation reduces Lsr2 binding to DNA, and ChIP-sequencing confirms increased DNA binding of a phosphoablative (T112A) Lsr2 mutant in *M. tuberculosis*. Altered expression of target genes in T112A Lsr2 compared to wild type Lsr2 *M. tuberculosis* offers further evidence that phosphorylation mediates expression of the Lsr2 regulon. Structural studies reveal increased dynamics of the Lsr2 DNA binding domain from a T112D phosphomimetic Lsr2 mutant, providing a molecular basis for decreased DNA binding by phosphorylated Lsr2. Our findings suggest that, the essential protein kinase, PknB controls *M. tuberculosis* growth and adaptations to the changing host environment by phosphorylating the global transcriptional regulator Lsr2.

## INTRODUCTION

*Mycobacterium tuberculosis* is a slow-growing bacterium that can replicate in humans and cause tuberculosis. The pathogen is able to rapidly shut-down its growth to persist in non-replicating states in infected individuals, which can be modelled in the laboratory^1^. *M. tuberculosis* adaptation to stressful conditions is accompanied by dramatic changes in global protein phosphorylation but the importance of these modifications is poorly defined^2^. *M. tuberculosis* has eleven serine/threonine protein kinases which play significant roles in growth, virulence and metabolism^3^. In particular, protein kinase B (PknB) is indispensable for *M. tuberculosis* growth due to its critical role in the regulation of peptidoglycan biosynthesis^4,5^. It is also important for *M. tuberculosis* survival in hypoxic conditions and resuscitation during reaeration^6^. However, the molecular mechanism for PknB-mediated adaptation to hypoxia is unknown.

We have recently shown that PknB-depleted *M. tuberculosis* can grow in a sucrose magnesium medium (SMM)^5^. Comparative phosphoproteomics of PknB-producing and PknB-depleted mycobacteria revealed substantial changes in profiles of phosphorylated proteins. Specifically, the transcriptional regulators Lsr2 and EthR showed increased phosphorylation in PknB-producing compared with PknB-depleted mycobacteria, indicating that these proteins may be PknB substrates. EthR is a TetR family transcriptional regulator that controls expression of EthA, a monoxygenase responsible for activation of the prodrug ethionamide^7^. Protein kinase F was found to phosphorylate EthR *in vitro*, which negatively regulated EthR binding to DNA ^8^.

Lsr2 is a DNA binding protein that combines properties of a nucleoid associated protein^9^ and a global transcriptional regulator^10^. Lsr2 has over 1000 binding sites in *M. tuberculosis*^11,12^. The precise role of Lsr2 in mycobacterial biology remains unclear, nevertheless parallels may be drawn with similar proteins from other bacteria. Lsr2 represents the first example of an H-NS-like protein identified outside Gram-negative bacteria, moreover *lsr2* was able to complement the *hns* mutant in *Escherichia coli*^13^. Similar to the H-NS proteins, Lsr2 has been proposed to bind to the minor groove of DNA^14^ and to possess DNA bridging properties^15^. Deletion of *lsr2* in *M. tuberculosis* resulted in severe growth impairment on solid media, defects in persistence and adaptation to changing oxygen levels^10^. Additionally, Lsr2 has been shown to help protect DNA from reactive oxygen species, and over-expression of Lsr2 improved survival of mycobacteria treated with hydrogen peroxide^16^. However, deletion of *lsr2* did not alter survival of *M. tuberculosis* exposed to hydrogen peroxide or mitomycin C ^10^. Transcriptional profiling of an *lsr2* deletion mutant revealed global changes involved in cell wall remodelling, respiration and lipid biosynthesis^10^.

Here we show that recombinant PknB phosphorylated Lsr2 at T112, a site that we identified previously by a proteomic approach, implying that PknB is directly responsible for controlling phosphorylation at this site. We demonstrate that phosphorylation of Lsr2 at threonine 112 is important for *M. tuberculosis* growth and adaptation to hypoxic conditions. Our results show that phosphorylation of Lsr2, or phosphomimetic T112D mutation, reduce Lsr2 binding to DNA, which is explained by increased dynamics in the DNA binding domain of phosphorylated Lsr2. Chromatin immunoprecipitation and transcriptomics further confirm that the T112A phosphoablative mutant of Lsr2 has an increased number of Lsr2-DNA binding events and modified expression of genes involved in *M. tuberculosis* stress responses and virulence. Based on our data, we propose that PknB-mediated phosphorylation controls Lsr2 binding to DNA in *M. tuberculosis*, providing a functional link between serine/threonine protein kinase signalling in replicating bacilli and regulatory networks that enable *M. tuberculosis* to survive dynamic environments during infection.

## RESULTS

### Transcriptome profiling of PknB-depleted *M. tuberculosis* reveals Lsr2-regulated gene expression signature

Two transcriptional regulators were significantly under-phosphorylated in PknB-depleted *M. tuberculosis*: Lsr2 and EthR^5^. To determine the resulting effect on gene expression, we compared the transcriptional profile of PknB-depleted versus PknB-producing *M. tuberculosis*, using the same pristinamycin-inducible PknB conditional mutant of *M. tuberculosis* (*pknB*-CM)^17^ in sucrose-magnesium medium (SMM) to support growth^5^. In total 99 genes were differentially expressed: 65 were induced and 34 repressed by PknB-depletion (Table S1). Two functional classes were identified in the genes overexpressed by PknB depletion: regulatory proteins and proteins involved in lipid metabolism. The predicted transcriptional regulators were *csoR*, *rv1129c*, *rv1460*, *rv2017*, *rv2250c*, *rv3334*, *sigB, whiB3* and *whiB6* (Table S1). These transcription factors have been shown to regulate copper homeostasis (CsoR)^18^, iron-sulphur cluster biogenesis (Rv1460)^19^, cholesterol catabolism (Rv1129c/PrpR)^20^, the enduring hypoxic response (Rv3334)^21^, multiple stress responses (SigB)^22^, redox stress and complex lipid biosynthesis (WhiB3)^23^ and virulence factor expression (WhiB6)^24^. The transcriptional signature of PknB-depletion resembled features of intracellular growth, which significantly overlapped with *M. tuberculosis* macrophage-derived RNA profiles from several studies as reflected by hypergeometric probability values: *6.7x10^−23^* ^25^; *7.34x10^−18^* ^26^; *3.57x10^−17^* ^27^ (Fig. 1). This profile was exemplified by induction of pathways involved in mycobactin synthesis (*mbtB*/*C*/*D*), complex lipid phthiocerol dimycocerosate (PDIM) biosynthesis (*fadD26*, *ppsA*/*B*/*C*/*D*), metabolism of alternative lipid carbon sources, the glyoxylate shunt (*icl*), the methylcitrate cycle (*prpD*/*C*, *prpR*) and triacylglycerol synthase (*tgs1*). The isoniazid inducible genes (*iniB*/*A*/*C*), responsive to cell wall stress and cell wall targeting drugs were also upregulated^28^. Four of the nine genes coding for alternative ribosomal proteins, *rpmB1*, *rpmB2*, *rpmG1*, *rpsN2*^29^ were induced by PknB depletion.

**Figure 1.**
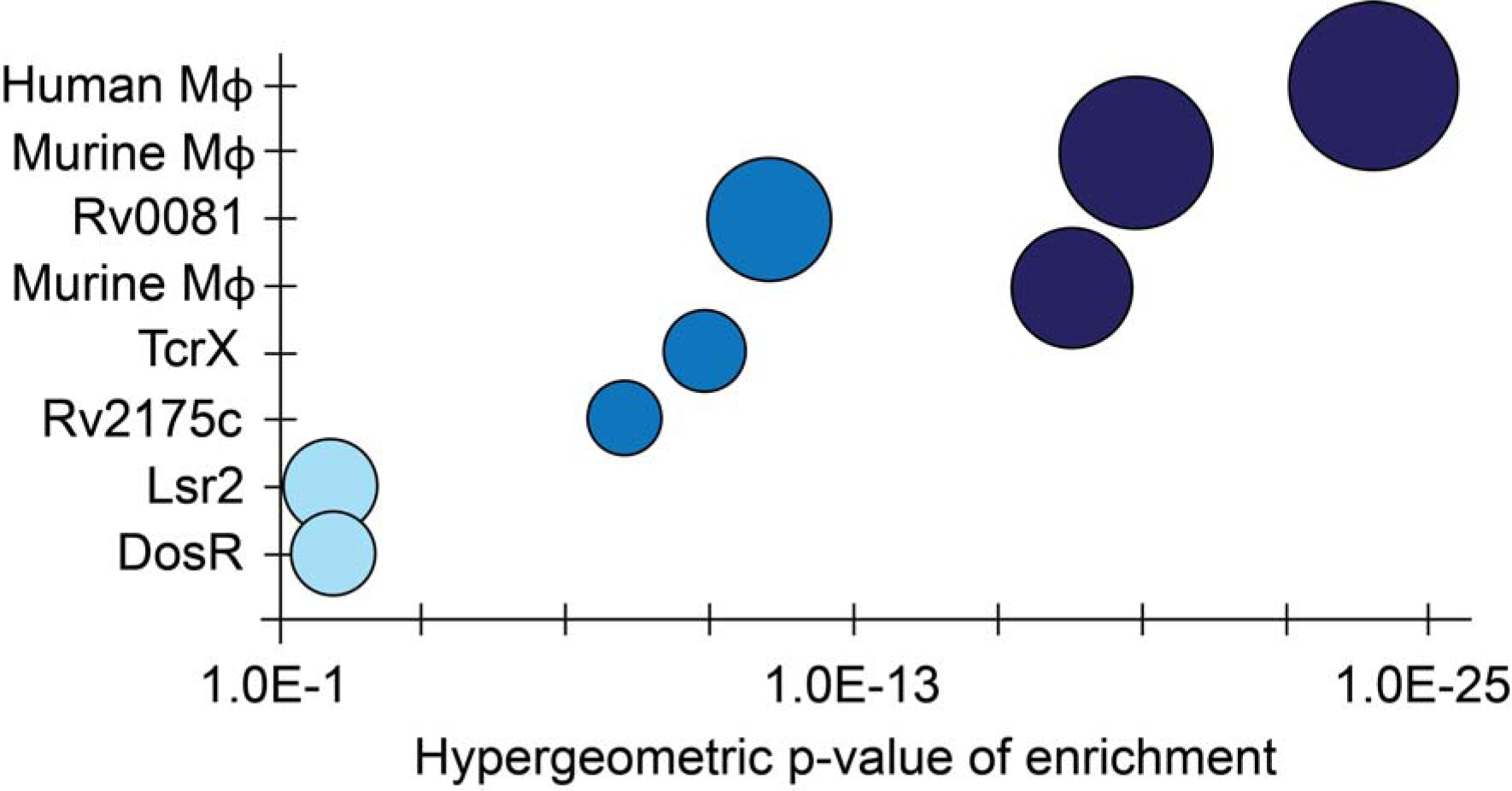
Transcriptional impact of PknB depletion in *M. tuberculosis*. The scheme highlights induced regulatory patterns from microarray (navy), transcription factor overexpression (blue), and ChIP-seq (light blue) studies referenced in the text. Enrichment p-value plotted on x-axis; bubble size proportional to number of overlapping genes in the signatures. P<0.02 is considered as statistically significant.

The 34 genes that were significantly repressed in PknB-depleted bacteria included *pknB* itself (6 fold down-regulated, as expected); *nuoA/B/C*, encoding subunits of NADH dehydrogenase I, which is part of the aerobic respiratory chain, and several genes involved in intermediary metabolism (Table S1). Overall these changes in replicating bacteria were comparable, in number of differentially expressed genes, to depletion of other regulators, for example DosR ^30^. This is in contrast to the large-scale changes in gene expression after treatment with an inhibitor of PknB and PknA that would likely impact *M. tuberculosis* viability ^31^. In summary, PknB depletion in replicating bacteria resulted in co-ordinated changes to the transcriptome, suggesting that PknB may control the induction of alternative gene regulatory pathways.

Application of the Transcription Factor Over-Expression (TFOE) output tool^32^ predicted Rv0081^33^, DosR^30^ and Lsr2^10^ as potential regulators of the observed gene expression patterns (Fig. 1). Rv0081 is a transcriptional hub which coordinates expression of over 50 transcription regulators controlling metabolic adaptation of *M. tuberculosis*^33^. However, to date there are no reports of Rv0081 regulation by phosphorylation. DosR has been shown to be phosphorylated by PknH^34^ but it was not under-phosphorylated during PknB depletion^5^. According to our data, PknB changed Lsr2 phosphorylation without impacting Lsr2 protein expression levels^5^, suggesting that the transcriptional changes between PknB-depleted and replete *M. tuberculosis* were likely caused by changes in PknB-mediated phosphorylation of Lsr2. We therefore investigated whether PknB could phosphorylate Lsr2 *in vitro*.

### PknB phosphorylates Lsr2 in vitro

Purified Lsr2 was mixed with the recombinant kinase domain of PknB and incubated at 37°C with or without ATP; phosphorylation was detected using western blotting with an anti-phosphothreonine antibody. Phosphorylated-Lsr2 was detected in the ATP-containing sample, demonstrating that PknB could indeed phosphorylate Lsr2 (Fig. 2A). Interestingly, phosphorylation resulted in a marked change in Lsr2 protein mobility on SDS-PAGE (Fig. 2B) and generated several bands, indicative of multiple phosphorylated isoforms. Mass-spectrometry confirmed the previously observed phosphosite on threonine 112^5^ and detected additional phosphorylations at threonine 8, threonine 22 and threonine 31 (Fig. 2C, D). The functional importance of these phosphorylation sites in Lsr2 was further investigated in complementation assays. We generated a panel of phosphoablative Lsr2 mutants in *M. tuberculosis* (Table S2) and investigated their phenotypes.

**Figure 2.**
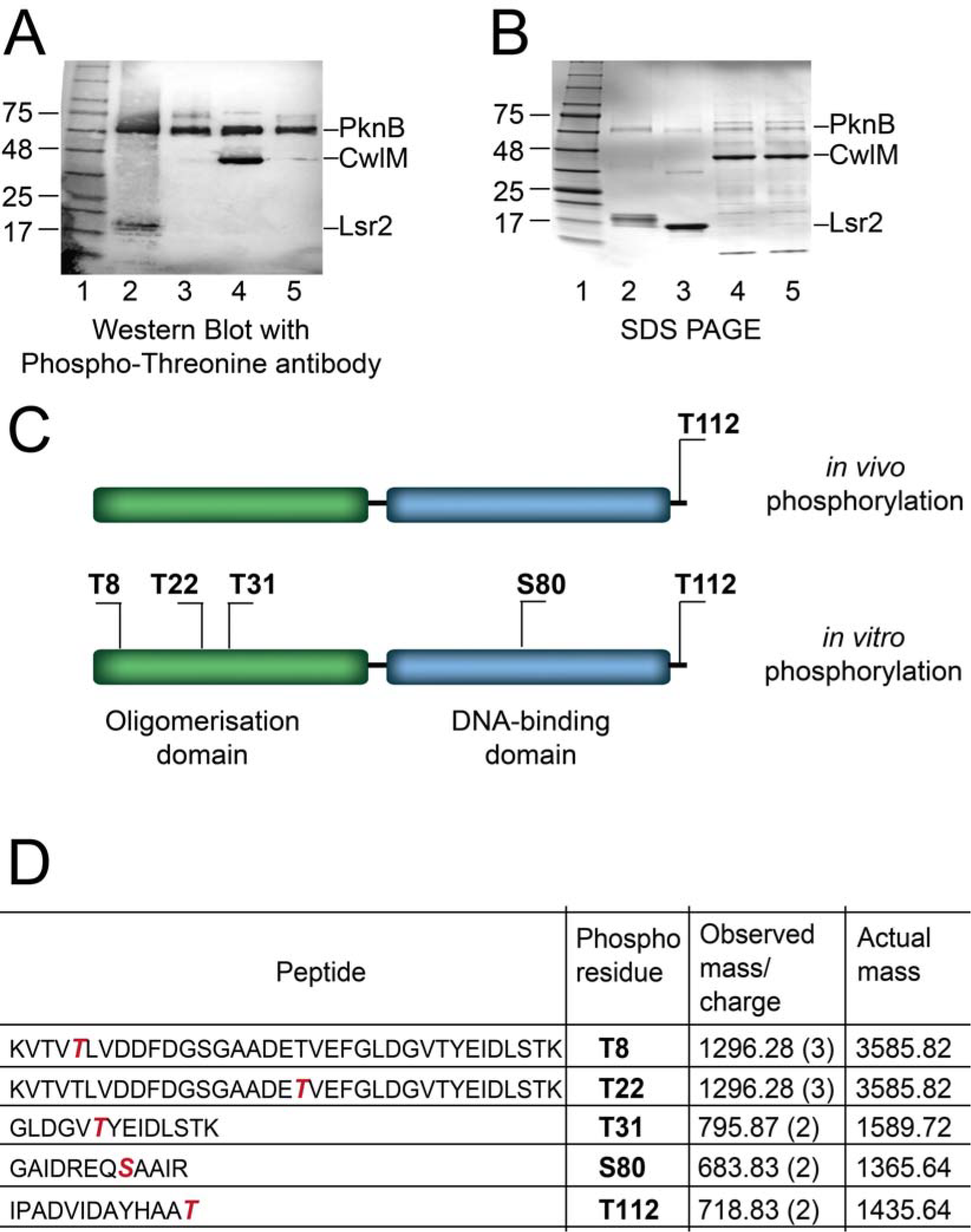
Identification of Lsr2 as a substrate of PknB. (A) Recombinant Lsr2 was phosphorylated by the recombinant enzymatic domain of PknB. Phosphorylated proteins were detected by western blot using a phospho-threonine antibody. Recombinant CwlM was used as positive control. 1-protein markers; 2 - Lsr2 incubated with PknB and ATP; 3 - Lsr2 incubated with PknB without ATP; 4 - CwlM incubated with PknB and ATP; 5 - CwlM incubated with PknB without ATP (B) SDS-PAGE revealed a shift in Lsr2 mobility upon phosphorylation (lanes identical to panel A). (C) Schematic presentation of phosphosites identified in phopshoproteomics studies (top)^5^, and *in vitro* (bottom). Lsr2 phosphorylation at T112-was 2.54-fold higher in PknB-producing *M. tuberculosis* compared to PknB-depleted *M. tuberculosis* (D) Phosphopeptides detected by mass-spectrometry; phosphorylated residues shown in red font.

### Phosphorylation of Lsr2 on threonine 112 is necessary for *M. tuberculosis* growth

*Lsr2* deletion significantly impaired *M. tuberculosis* growth on solid media and the reintroduction of *lsr2* on the pMV306 integrative plasmid (Δ*lsr2*_WT_) into the *lsr2* deletion mutant fully complemented this defect (Fig. 3A & Fig. S1), similarly to a previous study^10^. An Lsr2 deletion mutant containing the empty pMV306 plasmid (Δlsr2_pMV306_) had the same growth pattern as the *lsr2* deletion mutant (Fig. 3A), as expected. Lsr2 variants lacking the phosphorylation sites T8A, T22A and T31A individually restored bacterial growth and colony size to the same extent as wild type Lsr2 (Fig. S1). The T112A Lsr2 mutant (Δ*lsr2*_T112A_), however, failed to complement the Δlsr2 phenotype (Fig. 3A, Fig. S1). Replacement of threonine 112 with an aspartate to mimic phosphorylation (Δ*lsr2*_T112D_) complemented the Δlsr2 phenotype (Fig. 3A).

**Figure 3.**
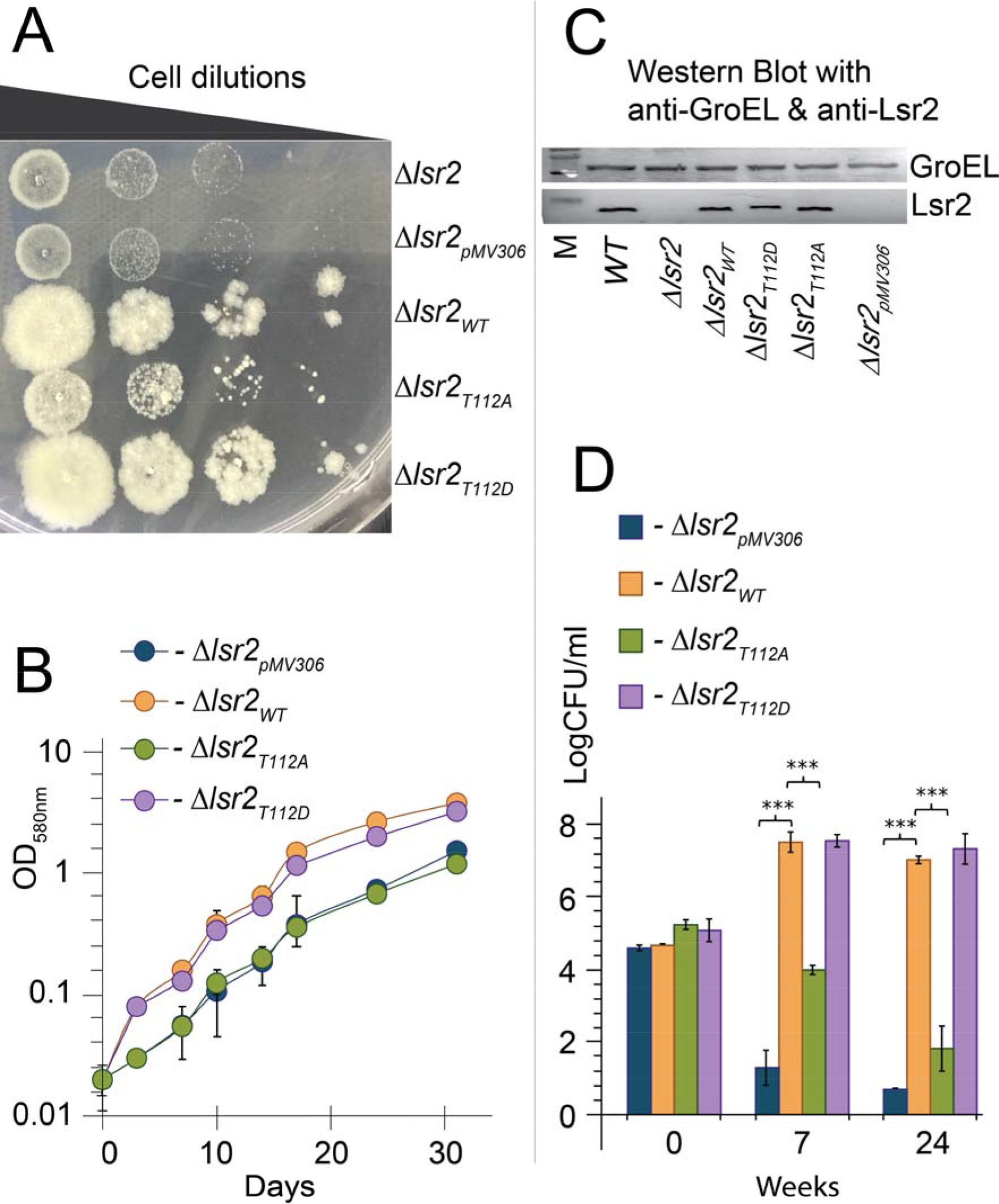
Phosphoablative T112A variant does not complement growth and survival defect of *lsr2* deletion mutant. Growth of *lsr2* deletion mutant expressing wild-type, T112A or T112D variants was compared with growth of the deletion mutant containing the empty vector. (A) 7H10 agar; (B) 7H9 liquid medium without shaking. Data presented as mean±SEM (N=6). (C) Expression of Lsr2 and Lsr2 variants from pMV306 was verified by Western blotting of *M. tuberculosis* lysates using an anti-Lsr2 antibody. Expression of GroEL was used as loading control. (D) T112A mutation impairs *M. tuberculosis* survival in the Wayne model of non-replicating persistence. *M. tuberculosis* Lsr2 mutants were incubated in sealed tubes with gentle mixing for up to 24 weeks. Data presented as mean±SEM (N=6). *** Statistically different in Δ*lsr2_pMV_* or Δ*lsr2_T112A_* compared with Δ*lsr2_WT_* and Δ*lsr2_T112D_* (p<0.001)

We next investigated whether T112 mutations of Lsr2 influenced *M. tuberculosis* growth in liquid culture. As Fig. 3B demonstrates, *Δlsr2*_pMV306_ and Δ*lsr2*_T112A_ had significant growth defects compared with Δ*lsr2*_WT_ and Δ*lsr2*_T112D_. Growth rates for *Δlsr2*_pMV306_ and Δ*lsr2*_T112A_ were 0.0157±0.03 h^−1^ and 0.0173±0.004 h^−1^, compared with 0.0414±0.004 h^−1^ and 0.0385±0.009 h^−1^ for Δ*lsr2*_WT_ and Δ*lsr2*_T112D_. Thus, the T112A mutant failed to complement the growth defect of Δ*lsr2* both in liquid and on solid media, suggesting that phosphorylation of Lsr2 at T112 is important for *M. tuberculosis* growth. Furthermore, western blot analysis using anti-Lsr2 antibody confirmed that wild type, T112A and T112D Lsr2 proteins were produced at similar levels, excluding the possibility that the Δ*lsr2*_T112A_ phenotype was caused by altered Lsr2 expression (Fig. 3C). The phosphomimetic T112D Lsr2 had lower mobility in SDS PAGE, similar to phosphorylated Lsr2 (Fig. 3C and Fig. 2B).

### Lsr2 phosphorylation at threonine 112 is crucial for survival in hypoxic conditions but not in prolonged stationary phase

We hypothesised that regulation of Lsr2 by PknB could account for the defects in survival of oxygen depletion seen when *pknB* and *lsr2* were perturbed^6,10^. Thus, we assessed survival of *M. tuberculosis* carrying Lsr2 variants using the Wayne model of non-replicating persistence in hypoxia^35^. The viable counts of Δ*lsr2*_pMV306_ dropped dramatically after 7 week-incubation compared to Δ*lsr2*_WT_ and Δ*lsr2*_T112D_ (Fig. 3D). Further incubation for 24 weeks resulted in near-complete loss of recoverable Δ*lsr2*_pMV306_ bacteria but did not significantly alter the survival of Δ*lsr2*_WT_ and Δ*lsr2*_T112D_. Interestingly, T112A Lsr2 could only partially complement the Δ*lsr2* survival defect, suggesting a requirement for T112 phosphorylation in microaerobic conditions. After 24 week-incubation, viable counts of Δ*lsr2*_pMV306_ and Δ*lsr2*_T112A_ were almost undetectable (Fig. 3D).

We also assessed the survival of *M. tuberculosis* expressing Lsr2 variants in late stationary phase of aerobically grown cultures by MPN and CFU counting. Δ*lsr2*_pMV306_ showed impaired survival compared to Δ*lsr2*_WT_, resulting in a 1.5 order of magnitude difference in viable counts. In this model, the survival of T112A or T112D variants was not significantly different from wild type Lsr2 (Fig. S1B). Our results suggest that Lsr2 phosphorylation of T112 may be specifically required during adaptation to hypoxic conditions but not for survival in prolonged stationary phase.

### Increased DNA binding in the T112A Lsr2 mutant of *M. tuberculosis* alters gene expression in *M. tuberculosis*

Phosphorylation of transcriptional regulators is known to influence their interaction with DNA ^8^. Lsr2 binds at multiple sites in the *M. tuberculosis* genome^14^. We hypothesised that Lsr2 phosphorylation might influence Lsr2-DNA interactions resulting in differential binding patterns in phosphoablative Lsr2 compared to wild type Lsr2 *M. tuberculosis*. We used a custom anti-Lsr2 antibody to define genome-wide Lsr2 binding patterns in *M. tuberculosis*. In control experiments, no DNA fragments were precipitated from Δ*lsr2* or Δ*lsr2*_pMV306_ mutants, confirming antibody specificity. We therefore determined the Lsr2 binding distribution from Δ*lsr2*_WT_, Δ*lsr2*_T112A_ and Δ*lsr2*_T112D_ in logarithmic phase *M. tuberculosis*. An average of 690 Lsr2 binding fragments were identified for wild type Lsr2 from three biological replicates (Δ*lsr2*_WT_), with a median length of 413 bp (Fig. S2, Table S3). Lsr2 binding was identified throughout the *M. tuberculosis* chromosome, with binding peaks often at intergenic spaces or with binding running through several open reading frames. The binding pattern was well-conserved across biological replicates (Fig. 4A, Fig. S2, Table S3). The putative regulon of Lsr2, including genes with an Lsr2 binding site within or immediately upstream of the coding sequence, was defined as 1178 genes (Fig. 4A, Table S3). These genes, potentially directly regulated by Lsr2, are consistent with previously identified Lsr2 binding patterns^14,36,12^ (Fig. S2). Interestingly, gene expression signatures associated with inactivation of Lsr2^10^, macrophage infection and acid-nitrosative stress were significantly enriched (hypergeometric probabilities 2.45x10^−27^, 3.02x10^−14^ and 8.59x10^−15^ respectively, Fig.S2), providing further evidence that Lsr2 may directly regulate gene expression.

**Figure 4.**
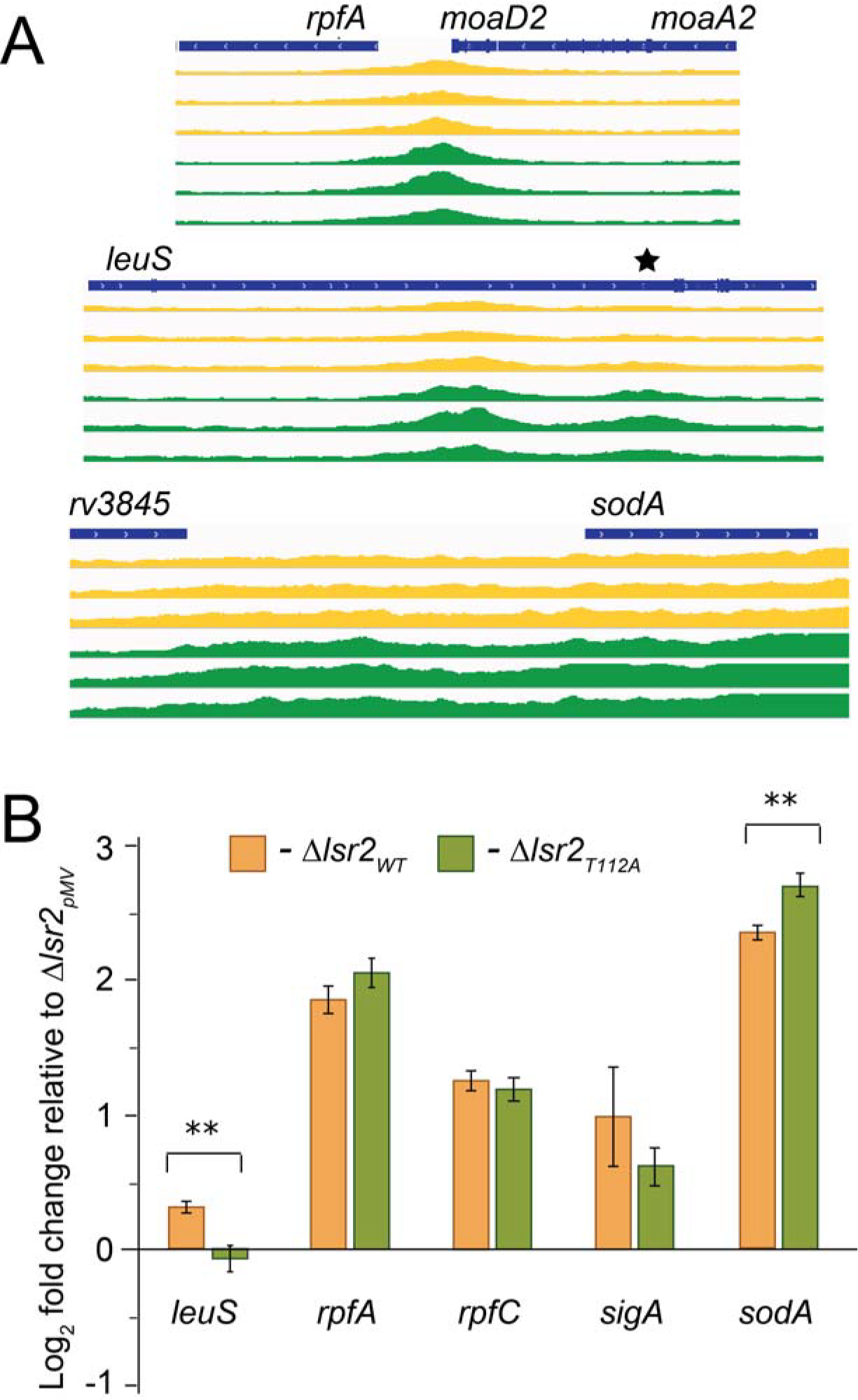
T112A mutation alters Lsr2 binding to DNA and gene expression patterns. **(A)** Representative plots describe Lsr2 binding upstream of *rpfA*, intragenic binding in *leuS*, or both intergenic and intragenic binding in *sodA*, showing greater Lsr2 binding in three biological replicates of phosphoablative Δ*lsr2_T112A_* (green) compared to Δ*lsr2_WT_* (yellow). A black asterisk marks the position of a Lsr2 binding site in *leuS*. Plots adapted from Integrative Genomics Viewer (IGV, ^63^). Binding patterns in Δ*lsr2_T112D_* were identical to those in Δ*lsr2_WT_* and not shown for clarity. (B) Expression of *leuS*, *rpfA*, *rpfC*, *sigA* and *sodA* relative to Δ*lsr2_pMV306_*measured by quantitative RT-PCR in Δ*lsr2_pMV306_*, Δ*lsr2_WT_*, Δ*lsr2_T112A_* and normalised to *16s rRNA and* Δ*lsr2_pMV306_*. Data presented as mean±SEM (N=6). **Statistically different in Δ*lsr2_T112A_* compared with Δ*lsr2_WT_* (p<0.01).

To establish whether phosphorylation changed the binding pattern of Lsr2 in *M. tuberculosis*, we compared phosphomimetic (*Δlsr2*_T112D_) and phosphoablative (*Δlsr2*_T112A_) to wild type Lsr2 (Δ*lsr2*_WT_). There were 226 binding events with significantly greater abundance in Δ*lsr2*_T112A_ in comparison to Δ*lsr2*_WT_ *M. tuberculosis* (Fig. S2, Fig. 4A). This potentially affected the expression of 94 genes (Table S4), including pathways involved in cell wall biosynthesis, lipid metabolism, PE/PPE protein synthesis and intermediary metabolism. No binding events were overrepresented in Δ*lsr2*_T112D_ compared to Δ*lsr2*_WT_. Thus, our results suggest that while phosphomimetic Lsr2 (Δ*lsr2*_T112D_) behaved similarly to wild type, phosphoablative Lsr2 (Δ*lsr2*_T112A_) bound DNA to a greater extent.

To probe whether altered binding of Lsr2 variants influenced gene expression we measured the expression of a panel of genes with predicted Lsr2 binding sites in Δ*lsr2_pMV306_*, Δ*lsr2*_WT_ and Δ*lsr2*_T112A_ genetic backgrounds. These genes included *leuS, rpfA*, *rpfC sigA*, and *sodA.* According to previously published data *rpfA*, *rpfC* and *sodA* were significantly down-regulated in the *lsr2* deletion mutant^10^, while expression of *sodA* and *leuS* were predicted to be differentially regulated in Δ*lsr2*_T112A_ in our ChIP-Seq analysis. Lastly, *sigA* encoding a housekeeping sigma factor, has a putative Lsr2 binding site in the promoter region^36^, which could influence its expression. Fig. 4B shows that 4 genes *rpfA, rpfC, sigA and sodA* were indeed up-regulated in Δ*lsr2*_WT_ compared with Δ*lsr2_pMV306_*. However, only expression levels of *leuS* and *sodA*, with divergent Lsr2 binding events, were statistically significantly different in Δ*lsr2*_T112A_ compared with Δ*lsr2*_WT_. Hence, our results confirm that the phosphorylation state of Lsr2 may influence gene expression patterns.

### PknB-mediated phosphorylation and T112D mutation reduce Lsr2 binding to DNA *in vitro*

We further investigated the impact of phosphorylation on Lsr2 binding to DNA *in vitro* using Electrophoretic Mobility Shift Assays (EMSA) with a range of DNA fragments containing putative Lsr2 binding sites. Lsr2 was able to shift all DNA fragments when added at concentrations above 1.9 μM; there was no significant difference in binding patterns between the DNA fragments (Fig. S3). For example, previously published data^36^ indicated a putative Lsr2 binding site in the *leuS* coding region (5’-AATTCGGC**AAAAT**CGGTAAG-3’ marked with an asterisk in Fig. 4A); this fragment was shifted in EMSA by Lsr2 (Fig. 5A). A DNA fragment containing a mutated Lsr2 binding site where the central AT rich region was replaced with a GC-rich sequence (5’-AACTCGGC**GAGGT**CGGTCAG-3’) was also shifted by Lsr2 (Fig. 5B). We tested other DNA fragments and all were shifted by Lsr2 regardless of their sequence, demonstrating that Lsr2 can bind non-specifically to DNA as previously suggested^28^ and in agreement with our ChIP-Seq analysis.

**Figure 5.**
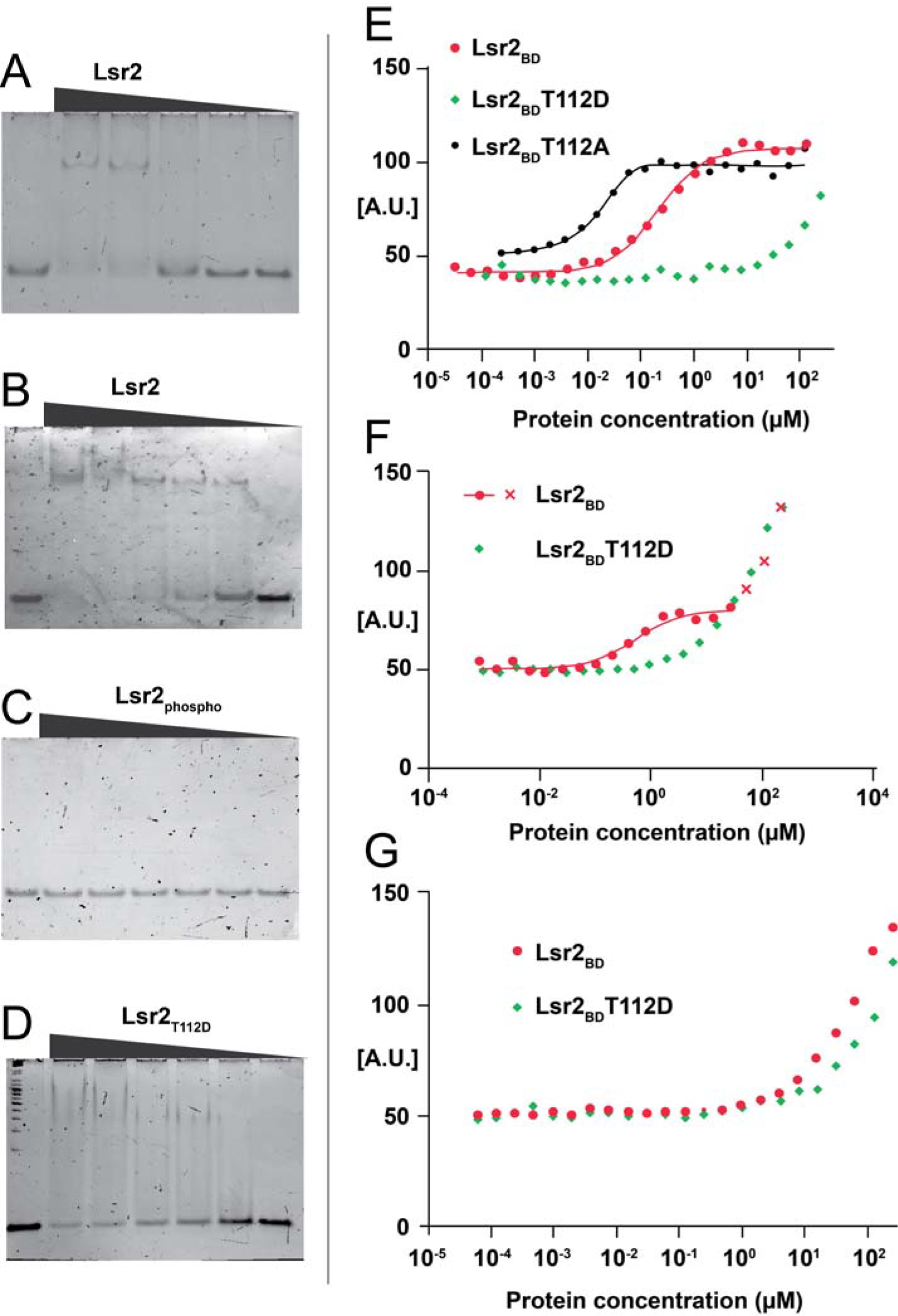
Phosphorylation or T112D mutation prevent Lsr2 binding to DNA. (A) Full-length Lsr2 was mixed with *leuS* fragment; (B) Full-length Lsr2 was mixed with the mutated *leuS*_MUT_ fragment; (C) Full-length Lsr2 was phosphorylated by PknB and mixed with *leuS* fragment; (D) Full-length T112D Lsr2 was mixed with *leuS* fragment. Lsr2 was added to DNA at a range of concentrations (0.95-7.6 μM). Representative results from three independent experiments shown. (E-G) Quantification of Lsr2_BD_ (DNA binding domain) interaction with DNA by fluorescence anisotropy. Titration of 5’ Alexa Fluor 488 DNA (4 nM) by Lsr2_BD_ WT (red), Lsr2_BD_ T112A (black) and Lsr2_BD_ T112D (green). (E) the previously identified DNA sequence (CGCGCATATATGCG); (F) *leuS* binding site (AATTCGGCAAAATCGGTAAG), red crosses designate points that were not used for Kd calculations; (G) mutated *leuS* binding site (AACTCGGCGAGGTCGGTCAG).

We next investigated the effect of Lsr2 phosphorylation on binding to the *leuS* site. As shown in Fig. 5C, phosphorylation of Lsr2 by PknB completely abolished binding (Fig. 5C). Similar results were obtained with other DNA fragments (Fig. S3). This *leuS* observation correlates with our ChIP-Seq finding (Fig. 4A) of a greater number of *leuS* binding events in phosphoablative compared to wild type Lsr2. Lsr2 T112D showed reduced DNA binding in EMSA experiments (Fig. 5D), suggesting that T112D mimicked phosphorylated Lsr2 to some extent but still showed weak binding to the fragments. Thus, full-length Lsr2 could bind DNA non-specifically and phosphorylation or T112D mutation abolished or reduced this binding *in vitro*. In H-NS proteins the oligomerisation domain is important for shaping the nucleoid, while the DNA binding domain regulates gene expression^37^. We therefore investigated whether phosphorylation of the DNA binding domain (Lsr2_BD_) influenced binding to DNA. The truncated protein Lsr2_BD_ was not suitable for EMSA, so we used a fluorescence anisotropy approach.

Previous studies demonstrated that Lsr2_BD_ mainly recognised AT-rich DNA sequences that formed a hook-like structure^14^. We measured Lsr2_BD_ and Lsr2_BD_T112D binding to an Alexa 488N labelled AT-rich DNA fragment (5’-CGCGCATATATGCG-3’), previously identified by Gordon and colleagues^14^ (Fig. 5E). At pH 7.5 the Kd value for Lsr2_BD_ was 0.21±0.06 µM, while the Kd value for Lsr2_BD_T112A was 0.02±0.008 µM and the Lsr2_BD_T112D showed no significant DNA binding (Kd could not be determined). Next, we tested binding of Lsr2_BD_ to the wild type and mutated *leuS* sequences where the central AT rich region was substituted by a GC-rich sequence (as described in the previous section). The binding of Lsr2_BD_ to the *leuS* sequence was characterized with a Kd of 0.4±0.12 µM (Fig. 5 F). No binding was observed for Lsr2_BD_ to the GC-mutated sequence (Fig. 5G). Lsr2_BD_T112D did not bind to either *leuS* sequences (Fig. 5F-G). These results show that T112D mutation, mimicking the Lsr2 phosphorylated state, reduced Lsr2_BD_T112D binding to DNA *in vitro*. Hence, phosphorylation controls both Lsr2 and Lsr2_BD_ binding to DNA.

### Phosphomimetic T112D mutation changes conformation entropy of the Lsr2 DNA binding domain

To elucidate a molecular basis for the reduced binding of phosphomimetic Lsr2 to DNA we compared nuclear magnetic resonance (NMR) structures of Lsr2_BD_WT and Lsr2_BD_T112D. In accordance with the previously published structure^11^, our data confirmed that Lsr2 _BD_ consists of two perpendicular α-helices (α1, residues 78–89; α2, residues 102–112) linked by a long loop (residues 90–101), and this organisation was preserved in Lsr2_BD_T112D (Table S5). However, T112D mutation resulted in a shorter α2 helix (Fig. S4), which ended with alanine 111 in Lsr2_BD_T112D compared with threonine 112 in Lsr2_BD_WT (Fig. 6A and 6B). Our data suggest that both proteins were in monomeric states and the N-terminal segment (residues 66-75) upstream of the DNA binding domain was disordered. We also found that in Lsr2_BD_ the methyl group of threonine 112 interacted with the tyrosine 108 and the hydroxyl group interacted with tryptophan 86 (Fig. 6C). These interactions did not form in the Lsr2_BD_T112D; however, they are unlikely to be important for DNA binding since T112A was able to bind DNA in our anisotropy experiments (Fig. 5E).

**Figure 6.**
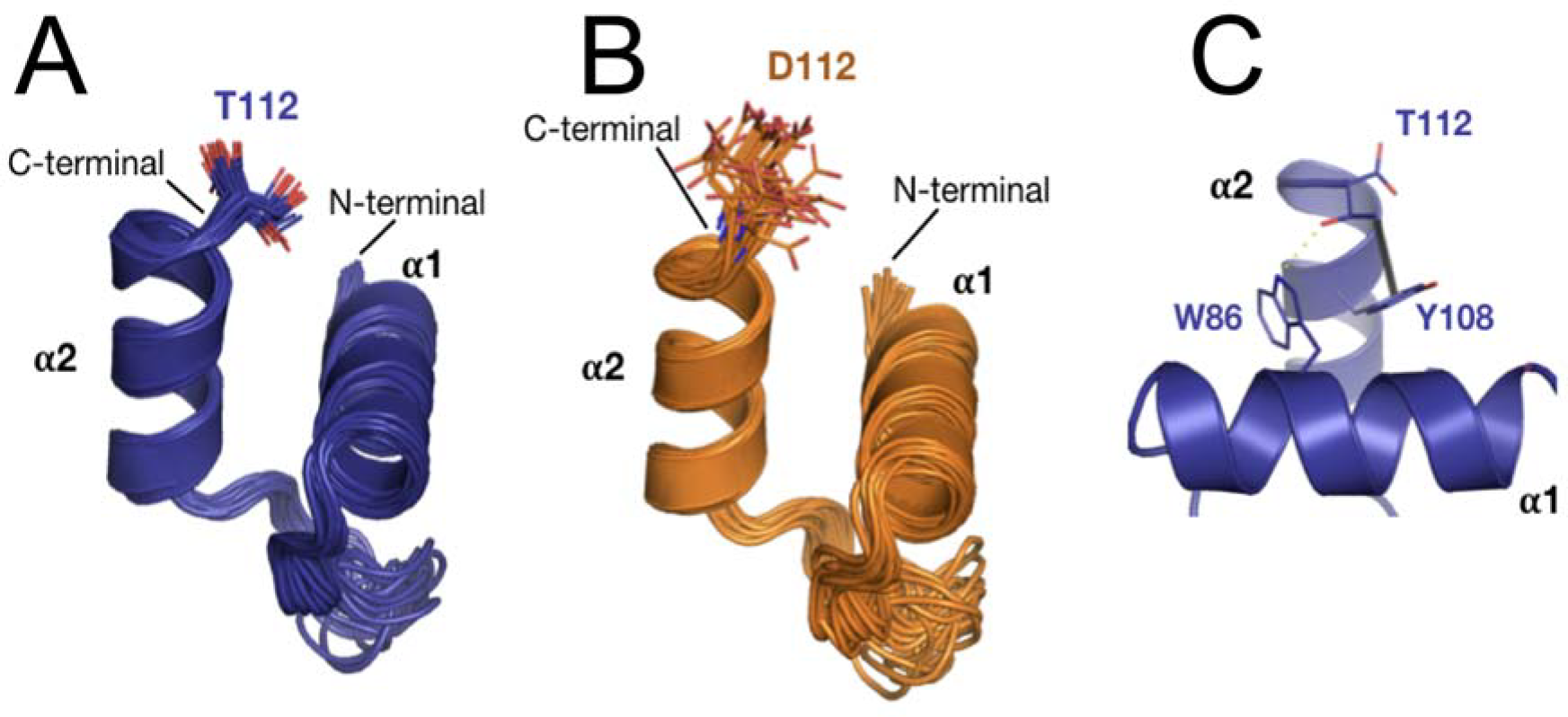
Solution structure of Lsr2_BD_ (A) and Lsr2_BD_T112D (B). Superimposition of the 20 best calculated structures in cartoon representation with the last residues represented as sticks (A and B). In Lsr2_BD_ the threonine T112 side chain interacts with both tyrosine Y108 (3.7 Å) and tryptophan W86 (3.0 Å), all represented in sticks (C).

^15^N heteronuclear NMR relaxation analysis was performed to assess the dynamic behaviour of the two proteins (Fig. S5). In both isoforms two α helices showed similar amplitudes for internal motion. However, in Lsr2_BD_ mainly the C-terminal helix was affected, while in Lsr2_BD_T112D the internal motion was extended to the N-terminal helix. A previous study on the catabolite activator protein (CAP) demonstrated that different protein mutants with the same structure of interaction interface displayed very different affinity for their target DNA^38^. The authors showed that changes of the binding affinity were linked to fast internal dynamics (conformation entropy). Similarly, the T112D mutation and, presumably, phosphorylation of threonine 112, resulted in a shorter α2 helix and a more mobile loop which increased the Lsr2_BD_ dynamics and impaired DNA binding.

## DISCUSSION

*M. tuberculosis* can subvert the immune system to survive in the host for many years. This remarkable ability is determined by mechanisms that allow *M. tuberculosis* to respond to multiple environments and adjust metabolic activity and cell division. One of these mechanisms is protein phosphorylation involving eleven serine/threonine kinases, including protein kinase B. PknB is expressed during active replication and is essential for growth in standard media ^17^. PknB has been implicated in the regulation of peptidoglycan biosynthesis^39^; its main substrate stimulates biosynthesis of peptidoglycan precursors ^4^and may facilitate their transport to the cell surface^5^. This regulation is essential for adjusting bacterial growth and synthesis of the cell wall. The external PASTA domain of PknB ^40, 41^ is essential for PknB-mediated signalling and its disruption results in bacterial death and alteration of antimicrobial susceptibility ^42, 5, 31^. Here we present data demonstrating that PknB also controls global gene expression via another substrate, the DNA-binding protein Lsr2. Transcriptomic analysis of PknB-depleted *M. tuberculosis* suggest that PknB may silence alternative pathways, which are not important for growth but may be critical for mediating stress responses and virulence. These include enzymes involved in alternative metabolic pathways, synthesis of complex lipids, regulators of stress responses and antimicrobial tolerance (Table S1).

Here we demonstrate that Lsr2 could be phosphorylated by PknB on several threonines but only the T112 was essential for growth and survival of *M. tuberculosis*. The phosphoablative T112A mutant could not complement the growth defect of the *lsr2* deletion mutant and was impaired in the Wayne hypoxia model. To investigate the molecular mechanism responsible for the growth defect in Δ*lsr2*_T112A_ we conducted ChIP-Seq analysis and compared DNA binding patterns in Δ*lsr2*_T112A_, Δ*lsr2*_T112D_ and Δ*lsr2*_WT_. In agreement with previously published data we detected multiple occurrences of DNA binding in all three Lsr2 backgrounds supporting both nucleoid shaping and gene regulatory functions of Lsr2. However, phosphoablative Lsr2 mutation T112A resulted in increased DNA binding, potentially directly affecting expression of 94 genes. Interestingly, most of these genes were not essential for growth, moreover deletion of some of these genes has been previously shown to be advantageous for growth^43^. The latter included genes of unknown function (*Rv0888*, *Rv1958*, *Rv1957*), PE/PPE genes (*Rv0878*, *Rv1983*), *aprA*, and genes controlling transport of PDIM (*drrB* and *drrC*). While products of these genes might be disadvantageous for growth *in vitro*, they likely play a critical role in adaptations to stress and virulence^27,44,45^.

We next investigated how phosphorylation affected Lsr2 binding to DNA. Our data demonstrate that PknB phosphorylation of Lsr2 *in vitro* completely abolished DNA binding, while the phosphomimetic mutation reduced Lsr2-DNA interaction. We have not investigated the effect of T8, T22 and T31 phosphorylations on DNA binding. Based on previously published data we hypothesise that phosphorylation of these sites in the oligomerisation domain might be important for controlling nucleoid shape and DNA bridging properties^37^. Our data suggest that phosphorylation of T112 in the DNA binding domain controls binding of Lsr2 to AT-rich DNA sequences which leads to alteration of gene expression. Our structural studies further confirm that phosphomimetic T112D mutation results in a shorter C-terminal helix and increased dynamics of the DNA binding domain, leading to impaired Lsr2-DNA binding.

Post-translational modifications are common mechanisms for regulation of DNA binding both in eukaryotes^46^and prokaryotes^47^. H-NS protein, a homologue of Lsr2 in *E. coli*, has been shown to be acetylated^48^ and phosphorylated^49^, however the precise function of these modifications remains to be characterised. Our data show that phosphorylation of Lsr2 is important for *M. tuberculosis* growth and this may be a key mechanism for controlling mycobacterial adaptations to permissive and non-permissive environments. Thus, PknB mediates two critical components of mycobacterial growth, peptidoglycan biosynthesis and gene expression of alternative pathways.

Based on our data we propose that PknB controls *M. tuberculosis* growth by phosphorylating Lsr2. Like other H-NS-like proteins Lsr2 plays a dual role in mycobacterial biology, it shapes and protects the nucleoid and it controls gene expression^9^. However, unlike H-NS proteins in Gram-negative bacteria which mainly silence the expression of foreign DNA^50,37^ Lsr2 regulates expression of genes that are essential for growth, virulence and adaptation^10^. Our data suggest that phosphorylation of T112 might be important for tuning gene expression during growth and the dynamic change between phosphorylated and non-phosphorylated Lsr2 may help to adjust transcriptional patterns according to growth conditions (Fig. 7A). Reduced T112 phosphorylation, for example during starvation, may increase Lsr2 binding and up-regulate pathways which are critical for *M. tuberculosis* survival under these conditions (Fig. 7B).

**Figure 7.**
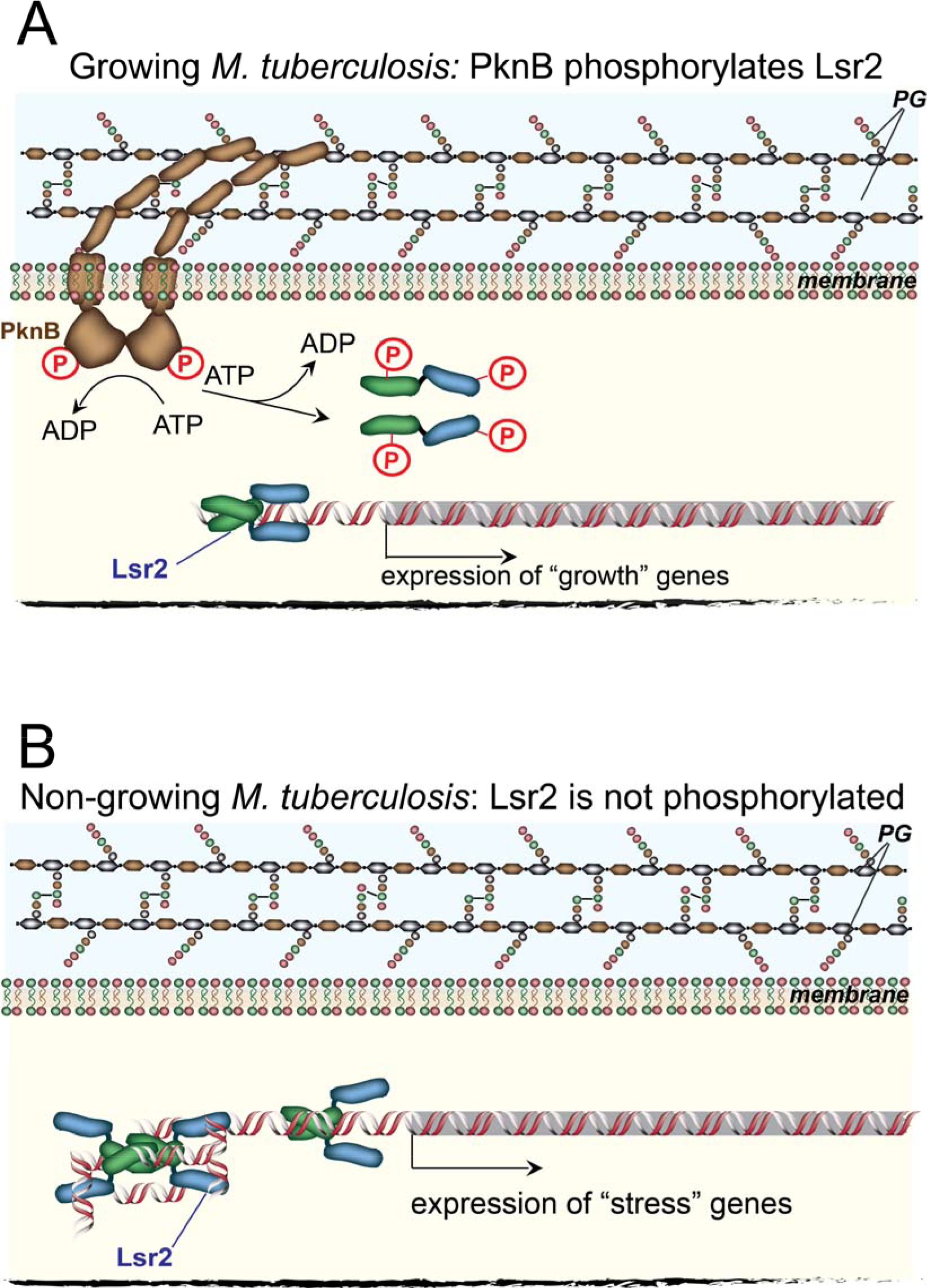
Lsr2 phosphorylation controls DNA binding and gene expression. (A) During growth PknB-mediated phosphorylation tunes gene expression to optimal levels and prevents activation of pathways disadvantageous for growth. Lsr2 is present as a mixture of phosphorylated and non-phosphorylated forms. The amount of phosphorylated Lsr2 is determined by activity of PknB, which is active during growth. (B) During non-permissive growth conditions, PknB is not produced and T112-nonphosphorylated Lsr2 binds DNA to a greater extent, switching on alternative transcriptional pathways and stress responses.

While there are many outstanding questions on the precise mechanisms of PknB-mediated regulation of gene expression and Lsr2 binding to DNA, our findings provide a functional link between serine/threonine protein kinase signalling and transcriptional regulatory pathways that enable *M. tuberculosis* to survive the varied environments encountered during infection.

## METHODS

### Organisms and media

*Mycobacterium tuberculosis* H37Rv was grown in Middlebrook 7H9 (Becton, Dickinson and Company) liquid media supplemented with 10% (v/v) Albumin-Dextrose Complex (ADC), 0.2% (v/v) glycerol and 0.1% (w/v) Tween 80 at 37°C with shaking. The *M. tuberculosis* H37Rv PknB conditional mutant (*pknB*-CM) was grown in sucrose magnesium medium (SMM) containing hygromycin with or without pristinamycin^5^. SMM comprised of 0.3M sucrose, 20mM MgSO_4_, 0.1% Tween 80 (w/v), 10% (v/v) ADC in standard 7H9 broth. Bacterial growth was measured by absorbance at 580nm, or by colony-forming unit (CFU) counting on 7H10 agar (Becton, Dickinson and Company), or by Most-Probable Number (NPN) counting using established protocols^51^; MPN counts were calculated using the MPN calculator program^52^. *Escherichia coli* OverExpress™ C41(DE3) and DH5α were grown in Lysogeny broth (LB). Antibiotics were used at the following concentrations (μg/ml): pristinamycin, 0.5; kanamycin, 50; hygromycin, 50; ampicillin, 50.

### Generation Lsr2 complementation mutants

The *Rv3597* (*lsr2*) coding sequence with additional 200 bp upstream region was amplified from the *M. tuberculosis* H37Rv genome using primers lsr2-pMV306F and lsr2-pMV306R and cloned into the integrative plasmid pMV306. Lsr2 *M. tuberculosis* variants were obtained using the GeneArt™ Site-Directed Mutagenesis System (Thermo Fisher Scientific) according to the manufacturer’s instructions. For generation of T112A and T112D mutants lsr2-pMV306RA and lsr2-pMV306RD primers were used. All constructs were sequenced before transformation into the previously described *lsr2* deletion mutant^10^. Bacterial strains generated in this study are shown in Table S2.

### Wayne model of non-replicating persistence

The model was setup as previously described^35^. CFU and MPN counts were determined at 0, 7 and 12-week time points. Statistical analysis was performed using one-way ANOVA (GraphPad Prism) where Δ*lsr2*_pMV_ or Δ*lsr2*_T112A_ data were compared to Δ*lsr2*_WT_ and Δ*lsr2*_T112D_ data.

### Transcriptomic analyses

*M. tuberculosis* H37Rv PknB conditional mutant (*pknB*-CM) was grown to OD_580nm_ ~0.7 in SMM with or without pristinamycin. *M. tuberculosis* RNA was extracted from three biological replicates using the GTC/Trizol method^53^. The RNA samples were purified and DNase-treated using RNeasy columns (Qiagen). *M. tuberculosis* RNA (2μg) was enzymatically-labelled with Cy3 fluorophore and hybridised to a *M. tuberculosis* complex microarray (ArrayExpress accession number A-BUGS-41) as previously described^54^. Significantly differentially expressed genes (Table S1) were identified using a moderated t-test (p-value <0.05 with Benjamini and Hochberg multiple testing correction), and fold change >1.8 in GeneSpring 14.5 (Agilent Technologies). Hypergeometric probability and TFOE analysis^32^ was used to identify significantly enriched signatures.

For quantitative RT-PCR, total RNA was isolated from triplicate *M. tuberculosis* cultures of Δ*lsr2*_T112A_, Δ*lsr2*_T112D_, Δ*lsr2*_WT_ and Δ*lsr2*_pMV_ as described above and cDNA generated using Superscript Reverse Transcriptase II and mycobacterial genome-directed primers^55^. Q-PCR was performed in a Corbett Rotor Gene 6000 real time thermocycler using Absolute QPCR SYBR Green mix and gene expression values were normalised to 16S rRNA expression. Statistical analysis was performed by one-way ANOVA where Δ*lsr2*_WT_ was compared to Δ*lsr2*_T112A_ and Δ*lsr2*_T112D_.

### ChIP-Seq analysis

DNA-Lsr2 interactions in Δ*lsr2*_T112A_, Δ*lsr2*_T112D_, Δ*lsr2*_WT_ were assayed using methods adapted from Minch *et al*^12^. Briefly, mid log-phase *M. tuberculosis* H37Rv cultures (OD_600_ 0.4-0.6) were crosslinked with 1% formaldehyde, followed by incubation with 125mM glycine for 5 minutes at 37°C. The cells were mechanically lysed and then sonicated to produce 200-500 bp fragments. Input control samples were taken for each genotype before antibody was added to assess antibody specificity. Samples were immunoprecipitated using a polyclonal anti-rabbit anti-Lsr2 antibody and protein-G agarose beads. The Lsr2 complexes were de-crosslinked by heating at 65°C overnight and proteins removed by treatment with proteinase K (10 mg/ml) for 2h at 55°C. The DNA samples were column-purified (Qiagen) and the quality of purified IP-Lsr2 DNA verified using the Qubit DNA HS quantification assay and Nanodrop spectrophotometer. Libraries were prepared and sequenced using Illumina HiSeq SE50, 20 million reads (Novogene, Hong Kong). Raw fastq files were aligned to the *M. tuberculosis* H37Rv (NC_000962.3) reference genome using bwa samse^56^. MACS2 (version 2.1.1.20160309)^57^ was used to compare each of the input controls to the immunoprecipitated samples, identifying Lsr2 binding sites (callpeak) using default parameters but including “-g 4.41e+06 –nomodel –extsize 147” (Table S3). Differential peaks comparing Δ*lsr2*_T112A_ to Δ*lsr2*_WT_ were then identified using “macs2 bdgdiff”.

### Generation and purification of recombinant proteins

The *M. tuberculosis* H37Rv *lsr2* coding sequence was amplified using Platinum Taq-HF polymerase (primers in Table S6) and cloned into pET15-TEV. After confirmation by sequencing, the constructs were transformed into *E. coli* OverExpress™ C41 (DE3) competent cells (strain details are summarised in Table S2).

For protein expression, *E. coli* cultures were grown in LB with ampicillin to mid-log phase (OD_600_ 0.6-0.8) at 37°C with shaking at 200rpm before adding 0.5mM isopropyl β-D-1-thiogalactopyranoside followed by incubation at 18°C overnight. The recombinant Lsr2 proteins were purified using immobilized-metal affinity chromatography (Ni-NTA agarose, Qiagen) and size exclusion chromatography. The recombinant catalytic domain of PknB was purified using Glutathione Sepharose 4B GST-tagged protein purification resin (GE Healthcare) as previously described^5^.

### Lsr2 phosphorylation *in vitro*

Purified recombinant Lsr2 (10 μM) was mixed with the recombinant catalytic domain of PknB (5 μM) in a kinase buffer (20 mM Tris–HCl, pH 8.0; 0.5 mM DTT; 10 mM MgCl_2_; 0.2 mM ATP) and incubated at 37°C for 1h. Phosphorylation was confirmed by western blot analysis using a phospho-threonine antibody (Cell Signaling Technology). Phosphorylated residues were identified in trypsin-digested proteins using a LTQ-Orbitrap-Velos mass spectrometer.

### Protein electrophoresis and western blotting

Proteins were separated on 4-20% gradient SERVA gels and transferred onto a nitrocellulose membrane using a Trans-Blot^®^ Turbo™ Transfer System (Bio-Rad). SIGMAFAST™ BCIP^®^/NBT or SignalFire™ Elite ECL Reagent were used to visualize proteins on C-DiGit Chemiluminescent Blot Scanner (LI-COR Biosciences). The following antibodies were used: custom polyclonal antibody raised against Lsr2 in rabbit (Gemini Biosciences); monoclonal murine anti-poly-histidine antibody (Sigma-Aldrich); phospho-threonine antibody (Cell Signaling Technology); monoclonal anti-*M. tuberculosis* GroEL2 (*Rv0440*), clone IT-70 (BEIResources); mouse anti-rabbit IgG antibody:alkaline phosphatase (Sigma-Aldrich; anti-mouse IgG (whole molecule:alkaline phosphatase antibody produced in rabbit (Sigma-Aldrich), and anti-rabbit IgG, HRP-linked antibody (Cell Signaling Technology).

### Electrophoretic mobility shift assay (EMSA)

EMSAs were carried out with DNA fragments amplified by PCR or annealed nucleotides (Table S6) as described previously^15^. Briefly, DNA (1.2 nM) was mixed with indicated amounts of Lsr2 in a total volume of 20µl reaction buffer containing (10 mM Tris–HCl, pH 7.5, 50 mM KCl, 1 mM DTT, 5 mM MgCl_2_, and 2.5% glycerol). The mixture was incubated for 30 min at room temperature followed by native polyacrylamide gel electrophoresis using 8% gels in 0.5 x Tris-Borate-EDTA buffer, pH 7.5 for 24 min at 120 V. The gels were stained with SYBR Safe DNA stain (Thermo Fisher Scientific) and visualised using a ChemiDoc system (Bio-Rad).

### Fluorescence anisotropy

Custom made 5’ Alexa Fluor 488 succinimidyl ester labelled oligonucleotide probe (sequence 5’-CGCATATATGCGCG-3) was purchased from Integrated DNA Technologies. Steady-state fluorescence anisotropy binding titrations were performed on a Tecan Saphire II microplate reader, using a 470nm LED for excitation and a monochromator set at 530nm (bandwidth 20nm) for emission.

### Determination of solution structures and dynamics of Lsr2_BD_ and Lsr2_BD_T112D

All ^1^H-^15^N double-resonance NMR experiments were performed at 20°C on Bruker Avance III spectrometers (700 or 800 MHz). NMR samples of 0.5mM ^15^N-labeled protein dissolved in 25mM sodium phosphate buffer (pH 6.8), 150mM NaCl with 10% D_2_O for the lock. ^1^H chemical shifts were directly referenced to the methyl resonance of DSS, while ^15^N chemical shifts were referenced indirectly to the absolute ^15^N/^1^H frequency ratio. All NMR spectra were processed with GIFA^58^ and analysed with CINDY (Padilla et *al.*, paper in preparation).

Chemical shift assignments were made using standard NOESY, TOCSY experiments performed on the ^15^N-labeled protein sample. NOE cross-peaks identified on 3D [^1^H, ^15^N] NOESY-HSQC (mixing time 160 ms) were assigned through automated NMR structure calculations with CYANA 2.1^59^. Backbone Φ and ϕ torsion angle constraints were obtained from a database search procedure on the basis of backbone (^15^N, HN, Hα) chemical shifts using TALOS+^60^.

For each protein, a total of 200 three-dimensional structures were generated using the torsion angle dynamics protocol of CYANA 2.1. The 20 best structures of each protein (based on the final target penalty function values) were minimized with CNS 1.2. All statistical parameters are summarised in Table S5. Relaxation rate constant measurements were performed on a 0.5mM protein sample, at 18.8 T (800 MHz). The pulse sequences used to determine ^15^N R_N_(N_z_) (R_1_), R_N_(N_xy_) (R_2_), and ^15^N{^1^H} NOE values were similar to those described^61^. The ^15^N longitudinal relaxation rates (R_N_(N_z_)) were obtained from 10 standard inversion-recovery experiments, with relaxation delays ranging from 18 ms to 1206 ms. The ^15^N transverse relaxation experiments (R_N_(N_xy_)) were obtained from 10 standard CPMG experiments, with relaxation delays ranging from 16 ms to 160 ms. Both series of experiments were acquired in two single interleaved matrices to ensure uniformity of the experimental conditions. Heteronuclear ^15^N{^1^H} NOE were determined from the ratio of two experiments, with and without saturation.

## Data availability

Fully annotated microarray data have been deposited in ArrayExpress; accession number E-MTAB-7627, https://www.ebi.ac.uk/arrayexpress.

ChIP-Seq datasets have been deposited in the European Nucleotide Archive (ENA); study accession number PRJEB31102, http://www.ebi.ac.uk/ena/data/view/PRJEB3110.

The coordinates for Lsr2 protein structures have been deposed in the PDB https://www.wwpdb.org; accession numbers PDB6QKP and PDB 6QKQ for Lsr2_BD_ and Lsr2_BD_T112D, respectively.

## ACKNOWLEDGEMENTS

The following reagents were obtained through BEI Resources, NIAID, NIH: Monoclonal Anti-*Mycobacterium tuberculosis* GroEL2 (Gene Rv0440), Clone IT70 (DCA4) (produced *in vitro*), NR-13657; Genomic DNA from *Mycobacterium tuberculosis*, Strain H37Rv, NR-48669. We acknowledge the Centre for Core Biotechnology Services at the University of Leicester for support with containment level 3 experiments and analysis of mycobacterial proteins. The project was supported by the UK Biotechnology and Biological Sciences Research Council grants BB/K000330/1 and BB/P001513/1 (GVM); the French Infrastructure for Integrated Structural Biology (FRISBI) ANR-10-INBS-05 grant (MCG), the Ministry of Higher Education and Scientific Research Iraq (KA), and the Wellcome Trust 204538/Z/16/Z (SJW).

## AUTHOR CONTRIBUTIONS

Conceptualization, OT, MCG, SJW and GVM; Methodology, CR, MW, AAW, HOH; Investigation KA, OT, PB, HJ, ADV, SJW, GVM; Analysis, OT, KA, HJ, PB, AAW, SJW, MCG, GVM; Resources, ILB, MIV, and AAW; Writing-Original Draft, GVM, SJW, MCG; Writing – Review and Editing, GVM, SJW, MCG, ILB, OT, HOH. Funding Acquisition, GVM, MCG, SJW, AK. Supervision, GVM, OT, SJW, HOH and MCG.

## SUPPLEMENTARY TABLES

Table S1. The transcriptional signature of PknB-depletion in *M. tuberculosis*, comparing pristinamycin-inducible PknB complemented strains in sucrose magnesium medium with and without pristinamycin.

A separate excel file

*Significantly differentially expressed genes were identified using a moderated t-test (p-value <0.05 with Benjamini and Hochberg multiple testing correction) and fold change >1.8 from three biological replicates. 65 genes were significantly induced and 34 genes were repressed by PknB-depletion in replicating *M. tuberculosis*.

**Table S3. The Lsr2 regulon.**

A separate excel file

**Putative regulons for wild type Lsr2 mapped to the *M. tuberculosis* H37Rv genome (Mycobrowser release 3 2018-06-05, ^62^). Individual ChIP-seq replicates are shown; the Lsr2 regulon was defined to have binding events in 2 of 3 biological replicates. Genes were included in the predicted regulon if the Lsr2 binding site was immediately upstream of, or within, the coding sequence of each gene. For Lsr2 intergenic binding between divergent genes, both genes were reported. For Lsr2 intergenic binding between convergent genes, neither gene was reported.

Table S4. Genes directly impacted by additional Lsr2 binding events in phosphoablative (T112A) Lsr2 *M. tuberculosis* compared to wild type Lsr2 *M. tuberculosis*.

A separate excel file

***Genes were included in the predicted regulon if the Lsr2 binding site was immediately upstream of, or within, the coding sequence of each gene. For Lsr2 intergenic binding between divergent genes, both genes were reported. For Lsr2 intergenic binding between convergent genes, neither gene was reported. Annotation from Mycobrowser (Release 3 2018-06-05, ^62^).

