## Supplementary material for "Protein kinase B controls *Mycobacterium tuberculosis* growth via phosphorylation of the global transcriptional regulator Lsr2": Figure S1-6, Table S2,S5, S6

[1]Leicester Tuberculosis Research Group, Department of Respiratory Sciences, University of Leicester, Leicester, LE2 9HN, UK; [2]Department of Basic Science, Faculty of Nursing, University of Kufa, P.O. Box 21, Kufa, Najaf Governorate, Najaf, Iraq; [3]Centre de Biochimie Structurale, CNRS, INSERM, University of Montpellier, 34090 Montpellier, France; [4]Wellcome Trust Brighton and Sussex Centre for Global Health Research, Brighton and Sussex Medical School, University of Sussex, Brighton, BN1 9PX, UK; [5]Core Biotechnology Services, University of Leicester, University Road, Leicester, LE1 7RH, UK; [6]Department of Immunology and Microbiology, University of Colorado School of Medicine, Aurora, CO 80045, USA; [7]Institute for Infection and Immunity, St George's University of London, London, SW17 0RE, UK. [8]LISCB, Department of Molecular and Cell Biology, University of Leicester, University Road, Leicester, LE1 7RH, UK.

$Contributed equally to this work

---

Supplementary figures 1-6

Supplementary tables 2, 5, 6

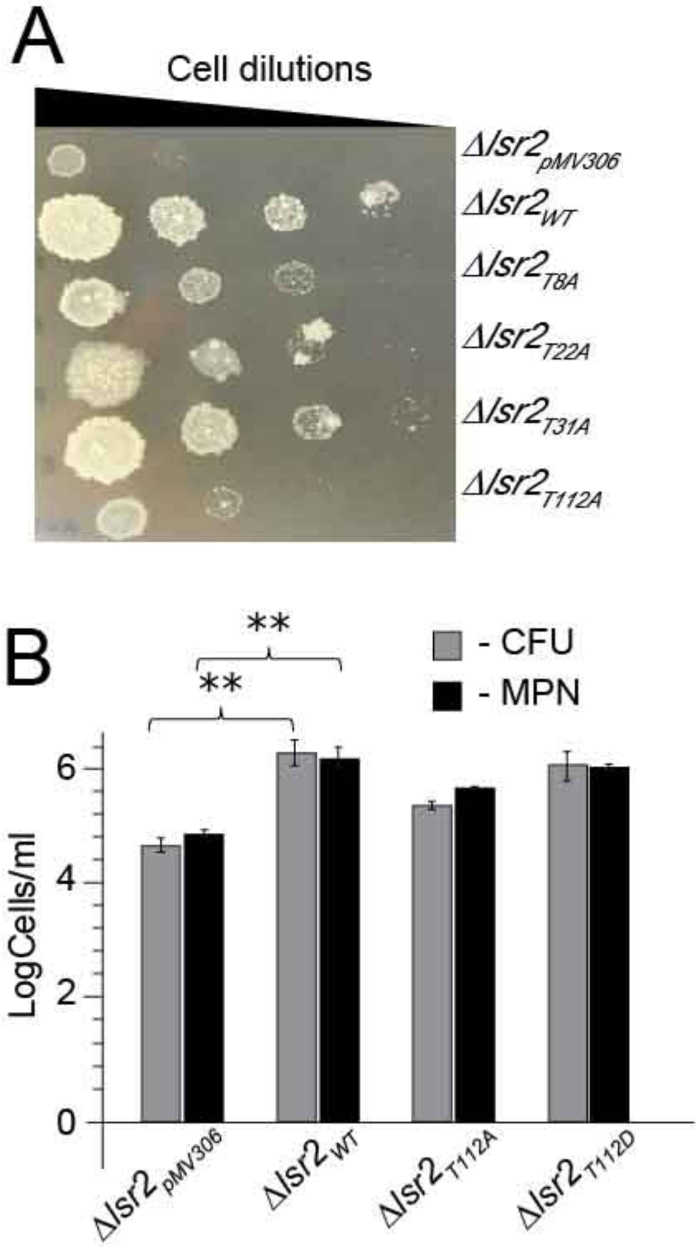

**Figure S1. Effect of Lsr2 mutations on growth on agar and survival in prolonged stationary phase.** (A) Lsr2 phosphoablative mutants were serially diluted and plated on 7H10 agar. (B) *M. tuberculosis* Lsr2 mutants were incubated with shaking for up to 42 weeks. Viable counts were determined by CFU and MPN counting. Represented as mean±SEM (N=6).

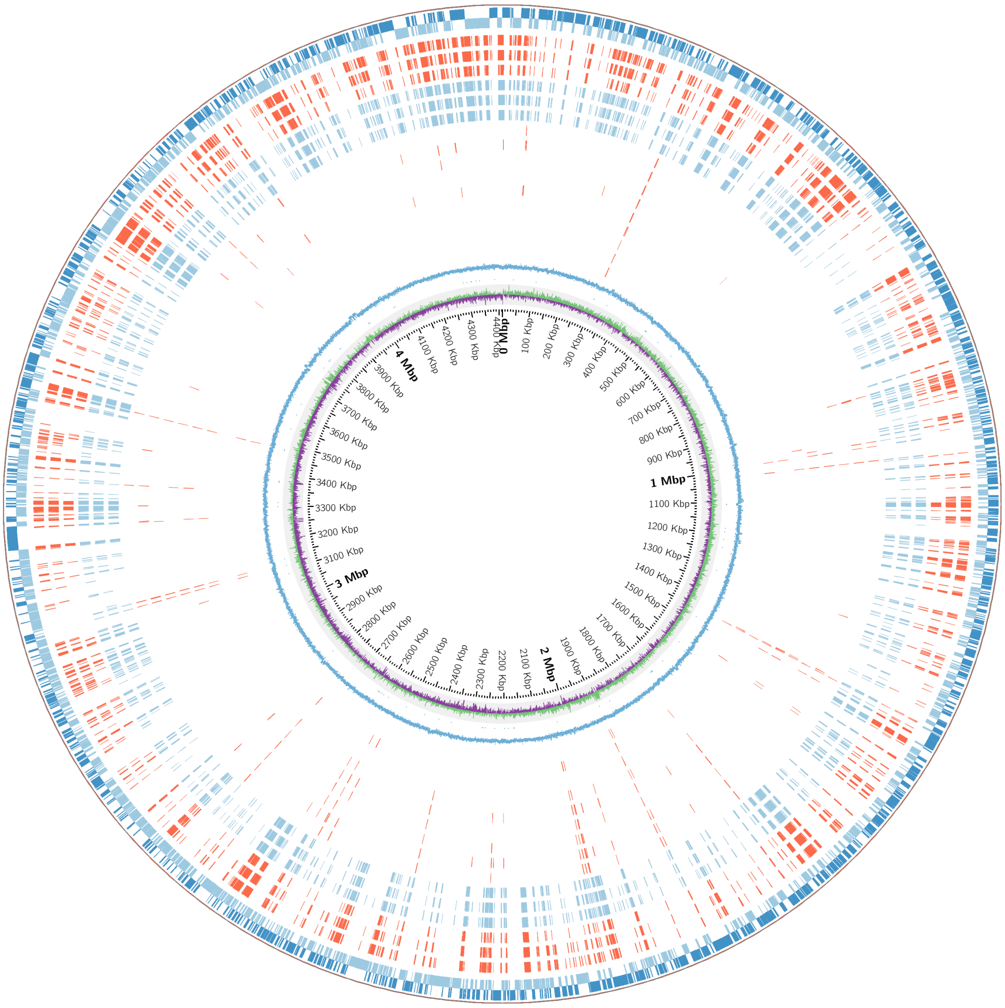

**Figure S2. The genome-wide binding pattern of Lsr2 in *M. tuberculosis*, showing the impact of phosphorylation on binding.** Moving from outside to inside, rings show the forward and reverse strands of the *M. tuberculosis* genome (blue), wild type Lsr2 binding events in 3 biological replicates (red), phosphoablative (T112A) Lsr2 binding sites in 3 biological replicates (blue), increase in abundance of Lsr2 binding in phosphoablative compared to wild type Lsr2 from pairwise comparison of biological replicates (red), GC% (blue), and GC skew (green/purple). These genes, potentially directly regulated by Lsr2, significantly overlapped with previously identified Lsr2 binding patterns from three independent studies, as indicated by hypergeometric p-values $1.36\times10^{-226}$ [1], $2.05\times10^{-190}$ [2], and $1.74\times10^{-84}$ [3], respectively. Interestingly, gene expression signatures associated with inactivation of Lsr2[4] were significantly enriched (hypergeometric probability $2.45\times10^{-27}$), providing further evidence that Lsr2 may directly regulate gene expression. Genes repressed in response to macrophage infection $3.02\times10^{-14}$ [5], sputum environment $8.70\times10^{-18}$ [6] and acid-nitrosative stress $8.59\times10^{-15}$ [7] also significantly overlapped with the predicted Lsr2 regulon, suggesting that Lsr2 may regulate *M. tuberculosis* adaptations to the changing environment.

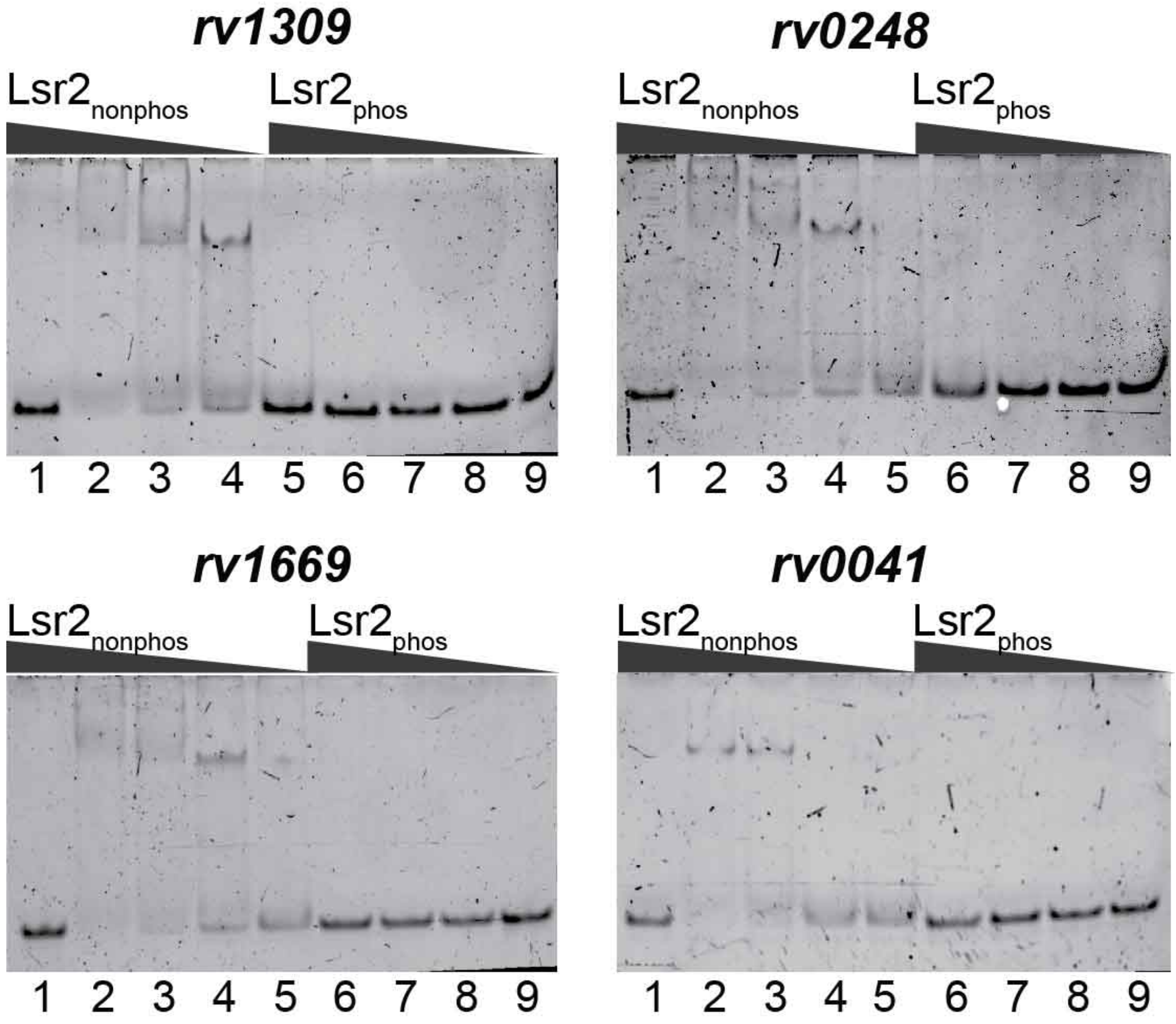

**Figure S3.** Lsr2 can shift various DNA fragments in EMSA. Lsr2 was mixed with annealed oligonucleotides containing putative binding sites within or upstream corresponding genes: *rv0041*, *rv0248*, *rv1306* and *rv1669.*

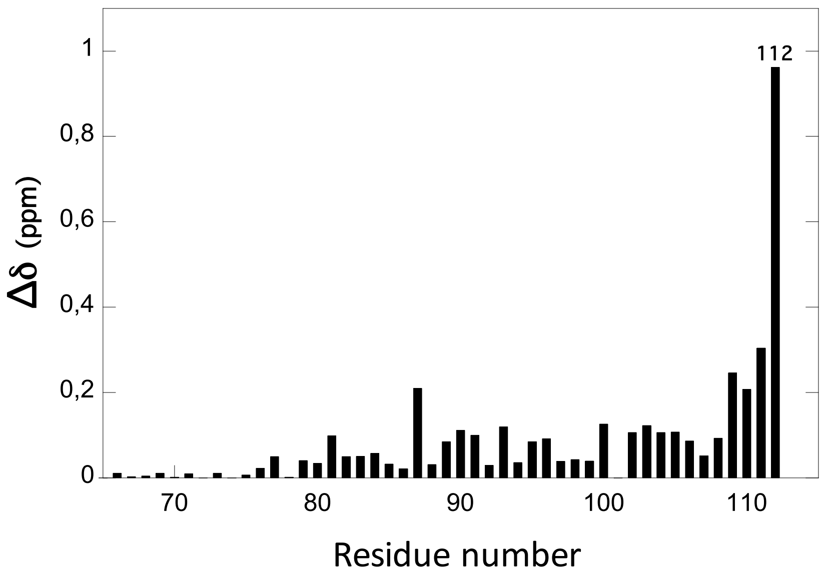

**Figure S4. Amide averaged chemical shift variations (Δ∂) as a function of the protein sequence.** Δ∂ have been calculated between the amide group resonances on $^{1}H$-$^{15}N$ HSQC spectra of $Lsr2_{BD}$ and $Lsr2_{BD}T112D$ with $\Delta\partial = [(\Delta\partial_H)^2 + (\Delta\partial_N \times (g_N/g_H))^2]^{0.5.}$

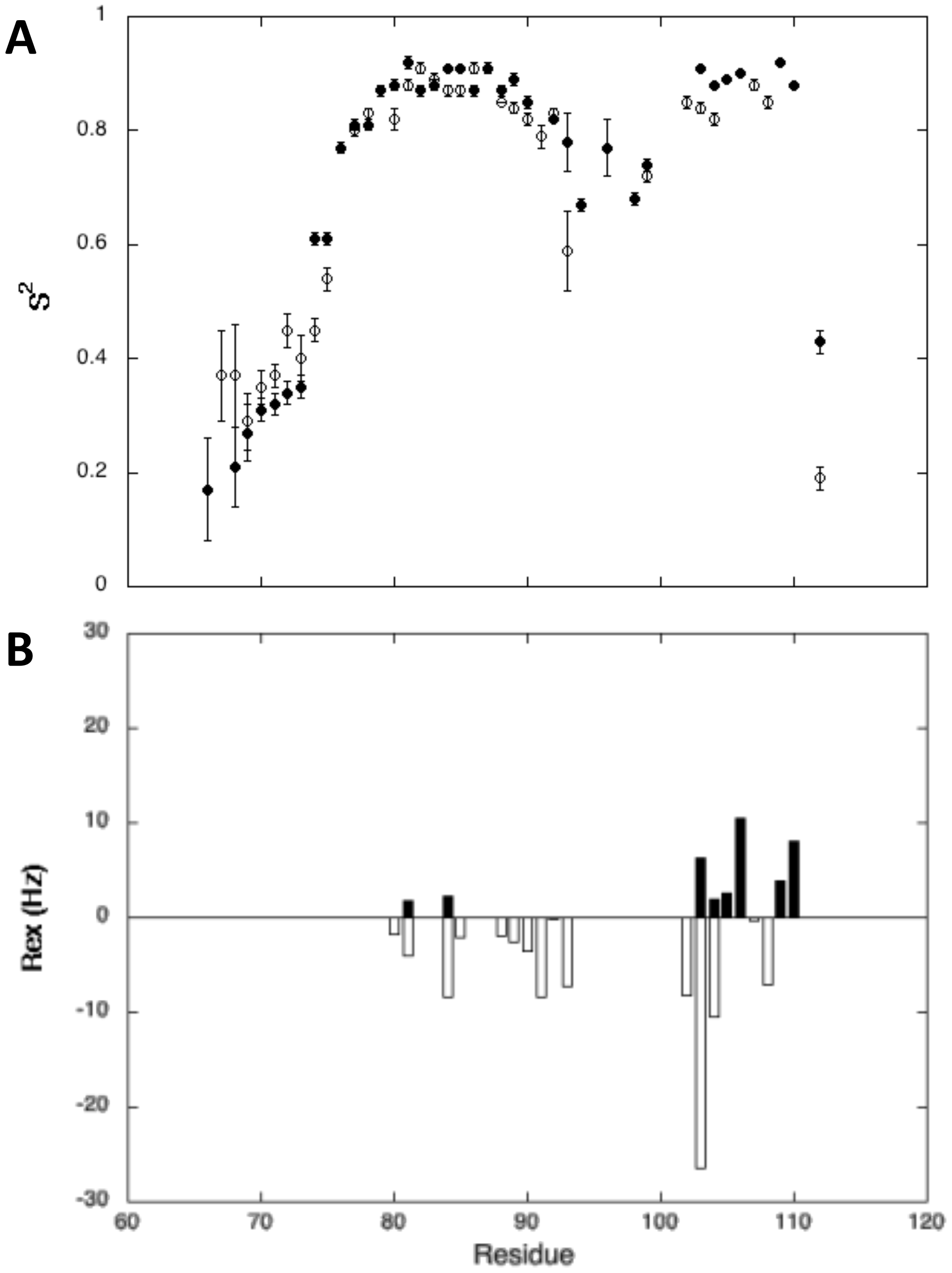

**Figure S5. Model-Free Dynamic Analysis of $Lsr2_{BD}$ WT and $Lsr2_{BD}$T112D.** (A) Generalized order parameters ($S^2$) values calculated for $Lsr2_{BD}$ (bold circles) and $Lsr2_{BD}$T112D (open circles). (B) Exchange contributions measured for $Lsr2_{BD}$ (bold bars) and $Lsr2_{BD}$T112D (open bars). For the sake of clarity, $R_{ex}$ values are reported on a negative axis for $Lsr2_{BD}$T112D. The global correlation time (3.90 ± 0.2 ns) extracted from Lipari-Szabo "Model Free" analysis of the $^{15}$N heteronuclear relaxation data ($T_1$, $T_2$, nOe) recorded on $Lsr2_{BD}$WT and $Lsr2_{BD}$T112D was consistent with a monomeric state for both proteins in our experimental conditions.

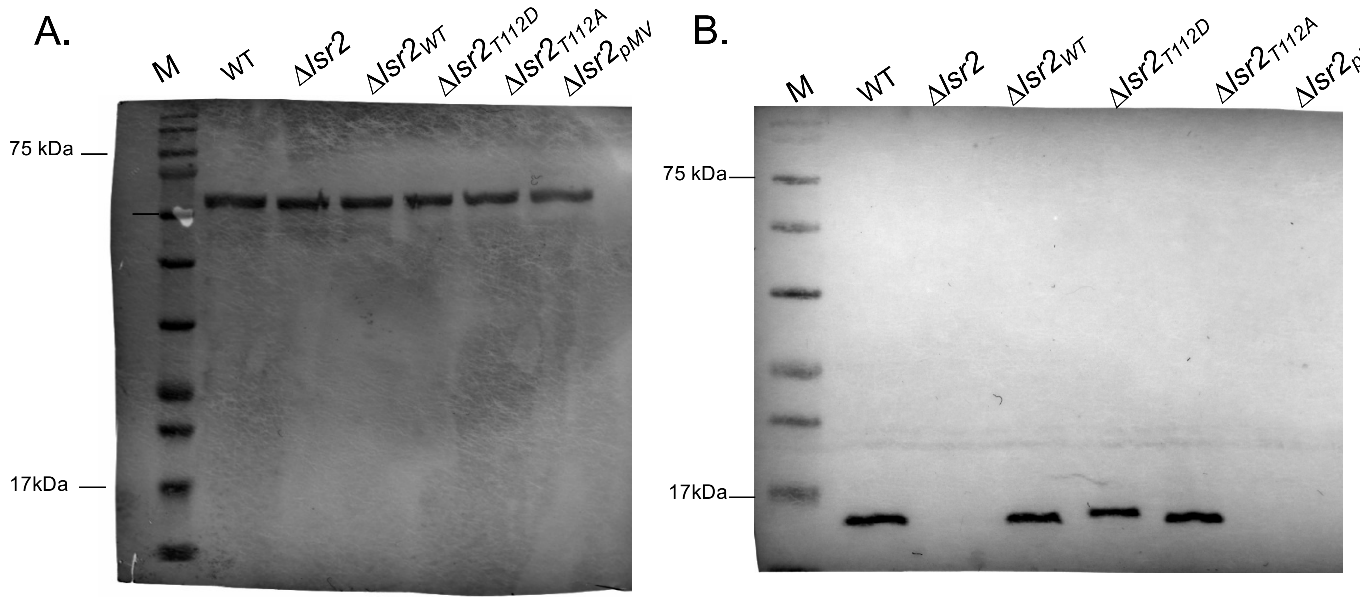

**Figure S6. Detection of GroEL (A) and Lsr2 (B) in *M. tuberculosis* lysates.** Lysates were prepared from growing M. tuberculosis cultures as described in Methods. A monoclonal anti-*M. tuberculosis* GroEL (BEIResources) and a custom polyclonal antibody raised against Lsr2 in rabbit were used for western blot analyses. M – protein markers, WT – wild type H37Rv strain. Three independent analyses were done, representative images shown.

**Table S2. Strains used and generated in the study.**

| Strain name | Strain description | Plasmid | Comments |
|---|---|---|---|
| ***Mycobacterium tuberculosis* H37Rv strains** | | | |
| *pknB*-CM | PknB conditional mutant | pAZI9479::*pknB* | PknB depletion |
| Δ*lsr2* | Deletion mutant | None | Lsr2 mutant background |
| Δ*lsr2*$_{pMV}$ | Empty plasmid control | pMV306 | Δ*lsr2* complementation |
| Δ*lsr2*$_{WT}$ | Δ*lsr2* complemented with wild type *lsr2* | pMV306::*lsr2* | Δ*lsr2* complementation |
| Δ*lsr2*$_{T8A}$ | T8A Lsr2 variant | pMV306::*lsr2T8A* | Δ*lsr2* complementation |
| Δ*lsr2*$_{T22A}$ | T22A Lsr2 variant | pMV306::*lsr2T22A* | Δ*lsr2* complementation |
| Δ*lsr2*$_{T31A}$ | T31A Lsr2 variant | pMV306::*lsrT31A* | Δ*lsr2* complementation |
| Δ*lsr2*$_{T112A}$ | T112A Lsr2 variant | pMV306::*lsr2T112A* | Δ*lsr2* complementation |
| Δ*lsr2*$_{T112D}$ | T112D Lsr2 variant | pMV306::*lsr2T112D* | Δ*lsr2* complementation |
| ***Escherichia coli* strains** | | | |
| C41(DE3) pET Lsr2 | 6xHis-Lsr2 expression strain | pET15bTEV::*lsr2* | Recombinant Lsr2 |
| C41 (DE3) pET Lsr2 T112D | 6xHis-Lsr2 T112D expression strain | pET15bTEV::*lsr2*$_{T112D}$ | Recombinant Lsr2 T112D |
| C41 (DE3) pET Lsr2$_{BD}$ | 6xHis-Lsr2 DNA binding domain expression strain | pET15bTEV::*lsr2*$_{BD}$ | Recombinant Lsr2$_{BD}$ |
| C41 (DE3) pET Lsr2$_{BD}$T112D | 6xHis-Lsr2 DNA binding domain T112D expression strain | pET15bTEV::*lsr2*$_{BD}$ *T112D* | Recombinant Lsr2$_{BD}$ T112D |
| C41 (DE3) pET Lsr2$_{BD}$ T112A | 6xHis-Lsr2 DNA binding domain T112A expression strain | pET15bTEV::*lsr2*$_{BD}$ *T112A* | Recombinant Lsr2$_{BD}$T112A |
| BL21 (DE3) pGEX PknB | GST-PknB kinase domain expression strain | pGEX::*pknB kinase* | Recombinant PknB kinase domain |

**Table S5: NMR and refinement statistics for LSR2-WT and LSR2-T112D peptide structures (LSR2-WT and LSR2-T112D, 0.5 mM, 25 mM NaPhosphate pH 6.8, 150 mM NaCl, 293 K)**

| | **LSR2-WT** | **LSR2-T112D** |
|---|---|---|
| **NMR distance and dihedral constraints** | | |
| Distance constraints | | |
| Total NOE | 616 | 482 |
| Intra-residue | 149 | 145 |
| Inter-residue | | |
| Sequential (\|*i* – *j*\| = 1) | 163 | 127 |
| Medium-range (\|*i* – *j*\| < 4) | 158 | 102 |
| Long-range (\|*i* – *j*\| > 5) | 146 | 108 |
| Hydrogen bonds | 40 | 30 |
| Total dihedral angle restraints | | |
| $\phi$ | 19 | 15 |
| $\psi$ | 19 | 15 |
| **Structure statistics** | | |
| Violations (mean and s.d.) | | |
| Max. distance constraint violation (Å) | 0.18 ± 0.05 | 0.15 ± 0.03 |
| Max. dihedral angle violation (°) | 1.21 ± 1.46 | 0.45 ± 0.44 |
| Deviations from idealized geometry | | |
| Bond lengths (Å) | 0.0098 ± 0.0005 | 0.0097 ± 0.0006 |
| Bond angles (°) | 1.1801 ± 0.0462 | 1.1952 ± 0.0663 |
| Impropers (°) | 1.2977 ± 0.1016 | 1.2639 ± 0.1166 |
| **Ramachandran plot (%)** | | |
| Most favoured region | 87.8 | 80.6 |
| Additionally allowed region | 10.4 | 16.5 |
| Generously allowed region | 1.2 | 1.4 |
| Disallowed region | 0.6 | 1.4 |
| **Average pairwise *r.m.s.* deviation** (Å)** | | |
| Backbone | 0.83 ± 0.31 | 1.13 ± 0.31 |
| Heavy | 1.72 ± 0.42 | 2.19 ± 0.46 |

** "Pairwise r.m.s. deviation calculated among 20 refined structures for residues 80-112."

**Table S6. Primers used in this study.**

| Primer name | Primer sequence (5'-3') | Comment |
|---|---|---|
| Lsr2testF | GTTGTGTCTGGATTGAGT | Lsr2 deletion confirmation |
| Lsr2testR | AAACCACCCAAGCGTTTC | Lsr2 deletion confirmation |
| pMV306lsr2F | CAC*GGTACC*GGAATGGGTATCGA | *Δlsr2* complementation |
| pMV306lsr2R | GACAAGCTTTCAGGTCGCCGCGT | *Δlsr2* complementation |
| pMV306lsr2AR | GACAAGCTTTCAGGCCGCCGCGT | *Δlsr2* complementation |
| pMV306lsr2DR | GACAAGCTTTCAGTCCGCCGCGT | *Δlsr2* complementation |
| T8AF | GCGAAGAAAGTAACCGTCGCCTTGGTCG ACGATTTCGAC | Lsr2 SDM |
| T8AR | GTCGAAATCGTCGACCAAGGCGACGGTT ACTTTCTTCGC | Lsr2 SDM |
| T22AF | TCGGGCGCCGCCGACGAAGCGGTCGAA TTCGGGCTTGAC | Lsr2 SDM |
| T22AR | GTCAAGCCCGAATTCGACCGCTTCGTCG GCGGCGCCCGA | Lsr2 SDM |
| T31AF | TTCGGGCTTGACGGGGTGGCCTATGAGA TCGACCTTTCC | Lsr2 SDM |
| T31AR | GGAAAGGTCGATCTCATAGGCCACCCCG TCAAGCCCGAA | Lsr2 SDM |
| Lsr-pETF | CAGCATATGGCGAAGAAAGTAACCGT | Recombinant Lsr2 |
| Lsr-pETR | CGACTCGAGTCAGGTCGCCGCGTGGT | Recombinant Lsr2 |
| Lsr-pETAR | CGACTCGAGTCAGGCCGCCGCGTGGT | Recombinant Lsr2 T112A |
| Lsr-pETDR | CGACTCGAGTCAGTCCGCCGCGTGGT | Recombinant Lsr2 T112D |
| pMV306F | TGGTATCTTTATAGTCCTGTC | pMV306 primer |
| pMV306R2 | TAGTTAACTACGTCGACATCGA | pMV306 primer |
| LeuSsiteF | AATTCGGCAAAATCGGTAAG | EMSA |
| LeuSsiteR | CTTACCGATTTTGCCGAATT | EMSA |
| Rv0248csiteF | GTCTTCCAATTGTCCTTCGG | EMSA |
| Rv0248csiteR | CCGAAGGACAATTGGAAGA | EMSA |
| Rv1306siteF | TGTGGCGATTTATCGTGCCG | EMSA |
| Rv1306siteR | CGGCACGATAAATCGCCACA | EMSA |
| Rv1669siteF | CTACCACATTAATCGGCATC | EMSA |
| Rv1669siteR | GATGCCGATTAATGTGGTAG | EMSA |
| LeuSsmutF1 | AACTCGGCGAGGTCGGTCAG | EMSA |
| LeuSsmutR1 | CTGACCGACCTCGCCGAGTT | EMSA |
| Rv0192F1 | AACACCGCGGTAAACATCGATGC | EMSA |
| Rv0192R1 | TACGTCGAGGAGTCCATGACCAC | EMSA |
| Rv3910F1 | TCAACTTGCTCAGCACCCACGGCAA | EMSA |
| Rv3910R1 | TGATCGATTCATGCGCGATCCGAAC | EMSA |
| leuSF1 | CATTTGGTCTACCGATCGTGGAAG | EMSA |
| LeusR1 | CTGTCATAGACGATCGGGAATGGT | EMSA |
| LeuSF2 | CACGTCAGCTCTCGCGAGCCTTAC | EMSA |
| LeuSR2 | GGTGTGCTCGTCGACGACCAAGC | EMSA |

| | | |
|---|---|---|
| LeuS F | AGCCAACGTCGTCAACTT | qRT-PCR |
| LeuSR | ATCCACTGAGTCCACCTGTA | qRT-PCR |
| SodAF | CACGTCAATCACACCATCTG | qRT-PCR |
| SodAR | GCACGGAACTTGTCGAAC | qRT-PCR |
| SigAF | GAG ATC GGC CAG GTC TAC GGC GTG | qRT-PCR |
| SigAR | CTG ACA TGG GGG CCC GCT ACG TTG | qRT-PCR |
| rpfAF | CTTGCAGTTCACTCAAAGCAC | qRT-PCR |
| rpfAR | CTCACCGACGGCAATCTG | qRT-PCR |
| rpfCF | AGCTGCCTCTCGGGAACAA | qRT-PCR |
| rpfCR | GACCACAGTGCGATCGGAAGG | qRT-PCR |
| 16s rRNAF | TCCGGGCCTTGTACACA | qRT-PCR |
| 16s rSRNAR | AACACCCGAAGCCAGTGG | qRT-PCR |
